# Follow-up availability and the robustness of immune transcriptomic recovery after myocardial infarction

**DOI:** 10.64898/2026.10.28.762142

**Authors:** Lingsen You, Guanzhao Zhang, Zisu Peng, Yujun Guo, Li Shen, Junbo Ge

**Affiliations:** Department of Cardiology, Zhongshan Hospital, Fudan University, Shanghai Institute of Cardiovascular Diseases, Shanghai 200032, China; National Clinical Research Center for Interventional Medicine, Shanghai 200032, China; Oriental Pan-Vascular Devices Innovation College, University of Shanghai for Science and Technology, Shanghai 200093, China; State Key Laboratory of Cardiovascular Diseases, Zhongshan Hospital, Fudan University, Shanghai 200032, China; NHC Key Laboratory of Ischemic Heart Diseases, Shanghai 200032, China; Key Laboratory of Viral Heart Diseases, Chinese Academy of Medical Sciences, Shanghai 200032, China; Institutes of Biomedical Sciences, Fudan University, Shanghai 200030, China; College of Biomedical Engineering, Fudan University, Shanghai 200438, China; Department of Cardiology, The Affiliated Hospital of Qingdao University, No. 16 Jiangsu Road, Qingdao 266000, Shandong, China

**Author notes:** Correspondence: Lingsen You; Junbo Ge. Lingsen You, Zisu Peng, and Yujun Guo contributed equally to this work.

**Keywords:** myocardial infarction, longitudinal transcriptomics, missing data, partial identification, inverse-probability weighting, reproducibility

## Abstract

**Background:** Longitudinal blood transcriptomes describe immune changes after myocardial infarction, but later assays do not necessarily represent the population sampled at admission. We examined whether recovery summaries depend on observable selection and assumptions about unavailable assays.

**Methods:** We reanalyzed official processed data from GSE59867, linking 111 patients with ST-segment elevation myocardial infarction across admission, discharge, one month, and six months. Seven fixed Hallmark programs were scored on a common 17,590-gene vocabulary using a bounded within-sample rank statistic. The primary endpoint was the six-month inflammatory-response score. We separated paired-completer change from the observed follow-up mean minus the entire baseline mean; fitted cross-fitted availability and outcome models; and evaluated full-support completion bounds and prespecified missing-mean scenarios. Artificial deletion tested estimators in the 83 observed six-month participants. GSE123342 provided unlinked marginal comparisons and explicitly conditional completion scenarios.

**Results:** GSE59867 contained 101, 95, and 83 follow-up assays. At six months, inflammatory-response change was −0.00893 among paired completers and −0.00999 for the marginal contrast (95% participant-bootstrap interval −0.01217 to −0.00783). Augmented inverse-probability weighting gave −0.01004 (−0.01556 to −0.00727). The full-support completion interval was [−0.16460, 0.08765], but eliminating the negative change required an unavailable-assay mean 4.42 observed-follow-up standard deviations above the observed mean. The primary direction remained negative throughout the prespecified ± 2-SD scenarios; interferon and coagulation directions were less stable. Under artificial outcome-dependent deletion, all evaluated estimators retained bias. The unlinked external cohort had 65 admission, 64 day-30, and 37 year-one MI assays; the year-one inflammatory contrast was −0.00459, with a conditional tipping shift of 1.70 SD.

**Conclusions:** The observed inflammatory-response rank-score decline was stable to baseline-score adjustment and the prespecified ± 2-observed-SD missing-mean scenarios, while unrestricted completion did not identify its direction. Robustness varied across programs and data settings. These findings concern relative transcriptomic scores and assay availability, not clinical recovery, survival, or intervention efficacy.

## 1 Introduction

Blood transcription changes during the acute response to myocardial infarction (MI) and subsequent recovery. Longitudinal peripheral blood mononuclear cell (PBMC) profiling has described inflammatory responses and potential markers of later heart failure, while whole-blood studies have related admission expression to ventricular dysfunction[1, 2]. Public expression repositories have enabled immune-atlas and recovery-signature analyses of these data[3, 4]. Such analyses can characterize the specimens that were measured. Attribution of their average change to the entire baseline population requires additional information about whose follow-up assays are available.

Multiscale immune forecasting also requires explicit observation populations. The Immune World Model preprint links cellular, tissue, and individual states in an action-conditioned model evaluated outside these MI cohorts; its therapeutic nominations remain prospectively testable hypotheses[5]. We examine a complementary requirement for longitudinal interpretation: which baseline participants are represented by available assays? We do not evaluate that model or its nominated intervention.

Three questions should be distinguished. Did measured individuals change? Did the composition of the observed population change? What would the average score have been if a defined assay had been obtained for every baseline participant? The first two can be examined when participant identifiers link visits. The third generally cannot be answered from observed assays alone. Assay nonavailability can reflect clinical loss to follow-up, death, planned subsampling, laboratory failure, consent restrictions, or incomplete deposition. The public record may not distinguish these processes. Treating all absence as biological deterioration, or assuming that complete cases represent the baseline cohort, adds unsupported information.

Missing-data methods provide a language for these distinctions. Inverse-probability weighting and augmented estimators can address selection explained by measured variables under explicit assumptions[6–8]. They cannot establish that outcomes unobserved at follow-up resemble those predicted from admission. Bounded outcomes also permit transparent worst-case completion intervals[9]. These intervals can be deliberately conservative: their width is determined by the missing fraction and outcome support. A useful empirical audit therefore needs both broad bounds and interpretable tipping points on the scale of measured variation. A narrow sampling interval around a completer estimate and a wide identification interval answer different questions.

The present study evaluates follow-up ascertainment in post-MI transcriptomic recovery. Here, transcriptomic recovery denotes temporal changes in the specified relative-expression scores after acute MI; it does not imply a validated normal target, a minimally important clinical difference, or clinical recovery. It does not introduce a new estimator or another diagnostic gene signature. Recent analyses already address MI diagnosis and recovery programs in these accessions[4, 10]. Our independent question concerns the population and assumptions to which a fixed recovery summary can be attributed. In contrast to recovery-signature construction, this analysis retains the complete authenticated 111-person baseline denominator and evaluates seven fixed programs whose membership was not selected using follow-up outcomes. It does not reconstruct or test the published recovery signature. We use a linked source cohort to quantify baseline composition, compare prespecified missing-at-random (MAR) estimators, and identify the missing-assay means required to reverse a conclusion. Artificial deletion supplies a known finite-data target against which estimator failures can be measured. A second, unlinked cohort illustrates which marginal summaries and conditional completion scenarios remain available when participant linkage is absent; it does not validate the source adjustment models.

## 2 Methods

### 2.1 Design, analysis chronology, and data roles

This is a secondary computational analysis of public processed microarrays deposited in GEO[11]. An internal specification fixed the primary score, primary visit, seven gene sets, estimators, sensitivity settings, and planned artificial-deletion mechanisms, fractions, and repetitions before scores or fitted models were computed. The exact weighted-deletion implementation differed from the frozen wording; this deviation was identified during post-analysis review and is documented below and in the dated implementation note. It was frozen on 27 October 2026 at 07:39:23 UTC (SHA-256 recorded in the reproducibility archive). Literature, sample metadata, availability counts, and prior reports of immune recovery had already been examined. The specification is not a preregistration and does not imply untouched external data. A metadata clarification recorded the lack of verified participant linkage and baseline-subset membership in GSE123342 before score analysis. Numerical summaries derived subsequently from the prespecified outputs do not create a new endpoint.

GSE59867 contains Affymetrix Human Gene 1.0 ST arrays (GPL6244) from a Warsaw STEMI study[1]. The original study enrolled consecutive patients treated with primary percutaneous revascularization during 2010–2013. This defines a reperfused STEMI setting; the present reanalysis does not isolate effects of revascularization or other treatment. The deposit contains 111 admission, 101 discharge, 95 one-month, and 83 six-month PBMC assays, plus 46 stable-coronary-artery-disease (CAD) controls. Explicit participant labels in sample titles link the 390 STEMI assays to 111 individuals. The primary denominator is these 111 participants. The controls are not part of the longitudinal estimand. The separate validation group reported in the original study is not added to the deposited expression cohort. Extreme-quartile heart-failure subgroup labels are not treated as full-cohort clinical outcomes.

GSE123342 contains whole-blood Human Transcriptome Array 2.0 measurements (GPL17586)[2]. Removing four explicitly labeled technical replicates leaves 65 admission MI, 64 day-30 MI, 37 year-one MI, and 22 CAD arrays. Public metadata do not supply authenticated participant keys. Although the source describes longitudinal sampling, the deposited record does not establish that every followup assay belongs to the specific 65 baseline individuals. We therefore use marginal descriptions and completion scenarios conditional on that additional subset assumption. We perform no pairing, participant-level weighting, or paired uncertainty estimation in this dataset. Clinical follow-up counts from the original larger enrollment group are not transferred to this array subset.

Two other matrices were downloaded during feasibility assessment but were not analyzed. GSE62646 was excluded because participant independence from GSE59867 was unresolved[12]. We do not pool cohorts, sum their arrays as independent patients, or describe six-month PBMC and year-one whole-blood assessments as matched follow-up horizons. Official accessions, sample inventories, technical-replicate mappings, download dates, file hashes, and conditions of use are retained in the archive and supplement.

### 2.2 Fixed transcriptomic measurement

We used the deposited processed expression values. Their original normalization may have used arrays jointly, including follow-up specimens; this analysis is not a reconstruction of prospectively isolated raw-data preprocessing. Probes with exactly one unambiguous annotated gene symbol were retained, and multiple probes for a gene were collapsed by their within-sample median. The common vocabulary comprised 17,590 genes represented on both platforms. This vocabulary depended on platform annotations and matrix probe identifiers, not on fitted effects or outcome selection. No pooled batch correction, external recalibration, or expression-driven gene selection was applied.

For each sample, genes were ranked from 1 to *G*, with average ranks for ties. For a fixed gene set *A* containing *k* retained genes, the score was

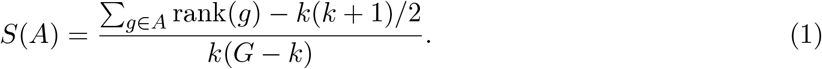

This statistic lies in [0, 1] and equals the fraction of gene-set/background gene pairs for which the set gene has higher expression, counting ties as one half. It is a relative rank-enrichment measure, related to single-sample rank-scoring approaches[13], rather than an absolute expression concentration or a validated measure of cell-intrinsic pathway activity. Fixed gene membership was taken from the public Hallmark collection[14]: inflammatory response (primary), TNF signaling via NF-*κ*B, interferon-*α* response, interferon-*γ* response, IL6/JAK/STAT3 signaling, complement, and coagulation. All passed the prespecified coverage requirement of at least 20 genes and 50% of published membership. Overlapping sets are not treated as independent biological replications.

### 2.3 Observed contrasts and hypothetical completion

Let *X*_*i*_ denote a baseline score, *Y*_*i*_ the corresponding follow-up score, and *R*_*i*_ indicate whether its assay is available among *n* baseline participants. Set *r* = *n*^−1^ ∑ _*i*_ *R*_*i*_, 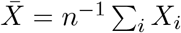, and 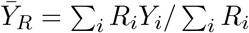. We distinguish the observed marginal contrast 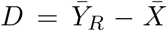 from the paired-completer contrast 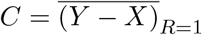 . Their difference is exactly

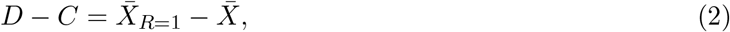

which measures observable baseline composition. Neither contrast alone estimates an unobserved whole-cohort follow-up mean without assumptions.

The completion target is the mean change if a defined follow-up score were available for every baseline participant. For an assumed mean *u* among unavailable assays,

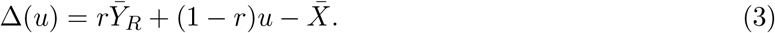

Under only the score support 0 *≤ u ≤* 1, the conditional completion interval is

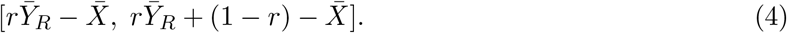

This is an identification interval conditional on the measured scores and roster, not a confidence interval or evidence that selection bias occurred. Its width is mechanically 1 *− r*. For *r <* 1, the zero-change tipping mean is 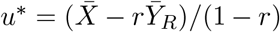; the required missing-versus-observed difference is *δ*^*\**^ = *−D/*(1 *− r*). We also report *δ*^*\**^ in observed follow-up standard-deviation (SD) units and evaluate the prespecified grid 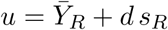, *d ∈* [−2, 2], clipped to [0, 1]. These scenarios do not estimate the missing outcomes or establish clinical plausibility. Observed ranges are contextual measurements, not guaranteed bounds on unobserved samples.

The hypothetical score must be defined for every person in the target. Death can make a later biological measurement undefined rather than simply missing[15]. Individual reasons for absence and survival are unavailable here. Consequently, the completion target is not a survivor-average effect, mortality-adjusted recovery, clinical prognosis, or treatment response. We do not solve truncation by death.

### 2.4 Observable selection and MAR estimators

At each source follow-up separately, assay availability was modeled from the seven admission scores using logistic regression with L2 regularization (*C* = 1) and no class balancing. Five stratified folds, fixed random seed 20261027, and training-fold standardization produced held-out probabilities. An intercept-only model supplied a predictive baseline. We report discrimination, Brier score, log loss, coarse calibration, and baseline-score differences; these assess availability prediction, not clinical prediction or the truth of MAR. Separate visit models accommodate intermittent missingness.

For observed follow-up assays, inverse held-out availability probabilities supplied normalized Hájek weights. Primary probabilities were clipped to [0.05, 0.95]; effective sample size and weighted baseline differences assessed weight behavior. A multi-output ridge model (*α* = 10), trained only on observed follow-up outcomes in each training fold with training-fold scaling, predicted follow-up scores for held-out individuals. Predictions were clipped to the known score support. G-computation subtracts 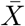 from the average held-out outcome prediction. The augmented inverse-probability-weighted (AIPW) estimator is

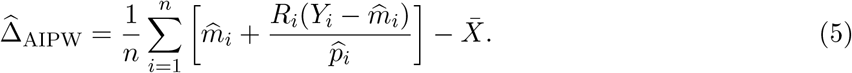

We do not truncate the resulting estimator. Weighting requires sufficiently correct availability probabilities, outcome regression requires a sufficiently correct conditional mean model, and the augmented estimator has established double-robust properties under MAR, positivity, and appropriate regularity conditions[7, 8]. Fixed clipping and regularization do not guarantee either model is correct. The standard double-robust guarantee is not automatically inherited by this clipped implementation: the propensity-correct branch additionally requires that clipping leave the relevant true probabilities unchanged; otherwise consistency requires a sufficiently correct outcome regression or additional conditions. The seven scores omit potentially relevant clinical, technical, and social determinants of assay availability.

Sensitivity analyses varied logistic *C* to 0.1 and 10, ridge *α* to 1 and 100, the number of folds to three and four, and probability clipping to none and [0.1, 0.9]. Settings were not selected according to their results. A 99-permutation availability-label analysis at six months reran the logistic pipeline while preserving the observed count. It is a pipeline diagnostic, not a test that missingness is ignorable.

### 2.5 Uncertainty and artificial deletion

Observed contrasts used 2,000 participant bootstrap resamples, with each person’s entire visit pattern kept together[16]. MAR estimates used 500 participant resamples with nuisance-model refitting; repeated draws of the same original participant remained in the same fold. Participant-bootstrap percentile 95% intervals summarize resampling variability of the observed-data estimators under an empirical participant-sampling model, conditional on deposited preprocessing, fixed gene sets, and the specified models. They are not confidence regions for the unidentified finite-cohort completion target or its support interval. Failures were logged and reported. These intervals do not incorporate uncertainty about the unknown missingness mechanism or provide external population guarantees. Secondary pathways and visits are descriptive; we did not conduct a family of confirmatory secondary significance tests.

Artificial deletion used the 83 actual six-month completers as a finite parent with known baseline and follow-up scores. Fifty masks at each planned fraction of 10%, 25%, and 40% removed 8, 21, and 33 assays, respectively. Selection was uniform, dependent on the baseline inflammatory score, or dependent on the follow-up inflammatory score. For the latter two, sampling without replacement used weights proportional to exp*{*2 clip(*z, −*2, 2)*}*, where *z* standardized the specified anchor score in the known parent. Higher-anchor assays were more likely to be removed. The frozen protocol described “rank-weighted sampling”, whereas the executed rule weighted standardized score values without an additional participant-rank transform. This was an implementation deviation: the mechanism categories, fractions, repetitions, and seed were retained, but the exact weighting formula was not fully prespecified. The original protocol was preserved unchanged, and the discrepancy was recorded after results were available. Each masked dataset was processed by the same availability and outcome models. Bias and root-mean-square error (RMSE) were calculated against the known parent mean change for unweighted, IPW, and AIPW estimates. Following the distinction between aims, data-generating mechanisms, estimands, methods, and performance measures in simulation reporting[17], the known parent target was held fixed. The 450 masks are repeated artificial perturbations of the same finite cohort. Methods are compared within each mask, and the masks do not represent new patients or independent clinical cohorts. These experiments expose behavior under specified mechanisms; they do not reveal the mechanism of actual missing assays.

### 2.6 Computational reproducibility and assistance

Python analysis code, exact dependencies, random seeds, source hashes, sample manifests, gene memberships, saved held-out predictions, numerical tables, and vector plotting code accompany the manuscript. The protocol hash is checked when the analysis runs. Independent computational checks recompute the rank statistic, patient counts, decomposition, bounds, and estimator formulas. Codex also assisted with literature triage, refinement of the research question, analysis specification and design, code construction, execution, numerical auditing, figure preparation, and drafting interpretations of the results and their limitations. No human–AI comparative study, autonomous scientific-agent performance experiment, reinforcement learning, or model self-improvement was conducted. The standardized declaration below is preserved as requested; release remains subject to author verification of the complete scientific and administrative record.

## 3 Results

### 3.1 The public record supports one linked longitudinal audit

The source matrix contains 436 arrays: 390 STEMI assays from 111 labeled participants and 46 CAD controls. Availability declined from 111 at admission to 101 at discharge, 95 at one month, and 83 at six months (Figure 1). Seventy-three participants had all four visits. Ten had an unavailable intermediate visit followed by an observed later visit, so a single monotone dropout-time model would misdescribe the deposit. The six-month absence fraction was 28/111 (25.2%). It describes missing deposited assays, not an established 25.2% rate of death or clinical loss to follow-up.

**Figure 1:**
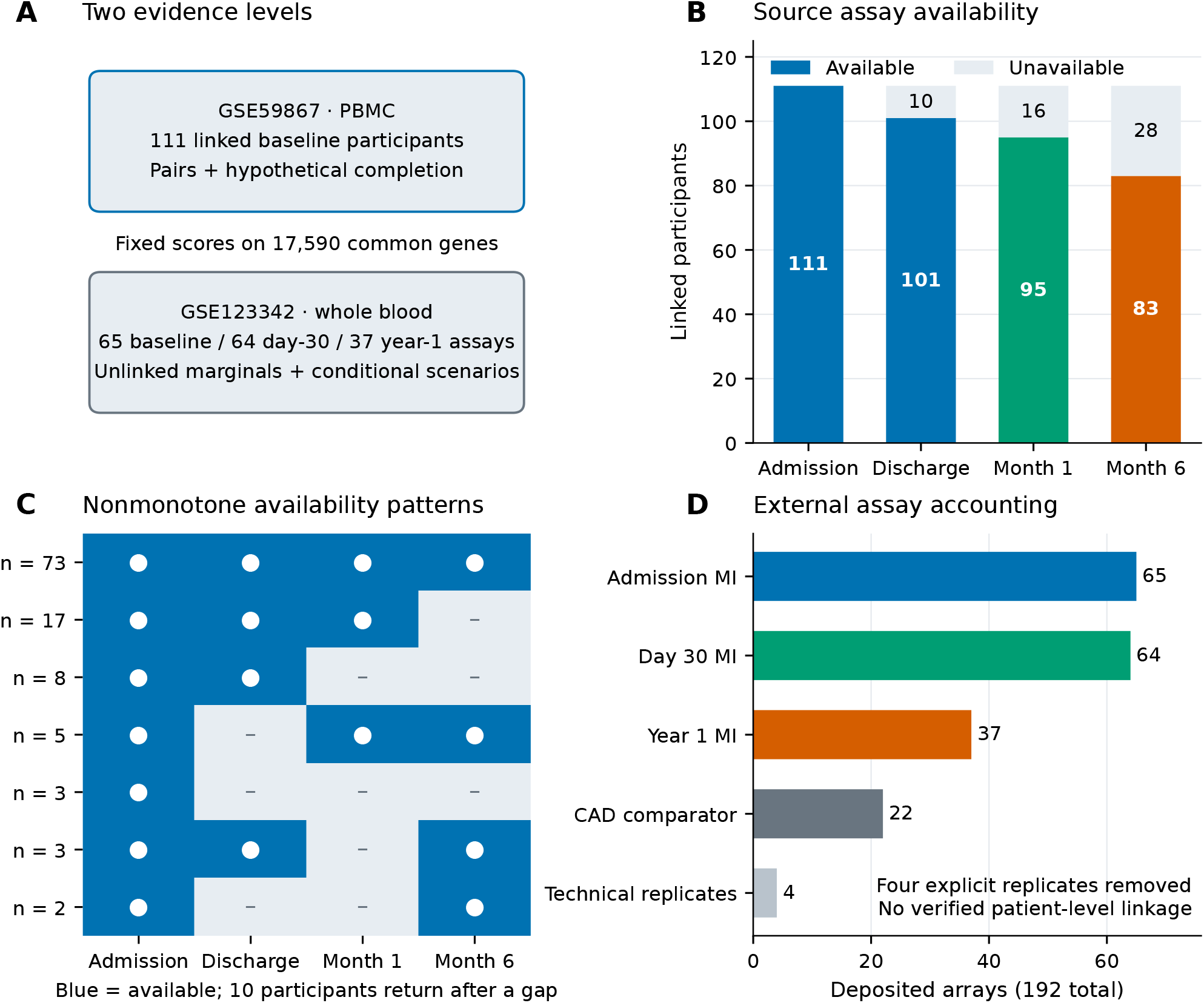
Evidence structure and follow-up assay availability. (A) The linked GSE59867 source series permits paired comparisons and assumption-dependent cohort-completion analyses. GSE123342 provides unlinked marginal comparisons; its completion scenarios additionally require unverified subset membership. PBMC, peripheral blood mononuclear cells. (B) Deposited source assays for the same 111 baseline STEMI participants; pale segments indicate unavailable assays, not documented deaths or withdrawals. (C) All observed visit-availability patterns, with their participant counts. Ten participants have a missing intermediate visit followed by an available assay. (D) Accounting of all 192 external deposited arrays. Four explicitly labeled technical replicates were removed; 188 arrays remain, including 22 CAD comparators not used in the MI temporal contrasts. Assay counts across unlinked external visits are not verified counts of retained participants. The two cohorts are not pooled into a claimed independent-patient total.

The measurement audit (Figure 8) documents coverage and overlap. A post-result descriptive plot of baseline score correlations was added to display dependence; it did not alter gene sets, estimators, or endpoints. The common annotation retained 191/200 inflammatory genes and 81–183 genes in each other program (83.5–95.5% of published membership across all seven). Every retained sample received all seven scores. The external dataset contained 192 arrays before removal of four declared technical replicates and 188 afterward. The 166 MI assays were distributed across 65 admission, 64 day-30, and 37 year-one records. These counts do not supply patient identities or certify a 37/65 patient-retention rate.

### 3.2 Inflammatory decline is visible in both paired and marginal summaries

Observed changes and their baseline-composition decomposition are shown in Figure 2. The source inflammatory-response mean was 0.62291 at admission and 0.61293 among the 83 available six-month assays. The marginal contrast was −0.00999 (95% bootstrap interval −0.01217 to −0.00783), compared with a paired-completer change of −0.00893 (−0.01124 to −0.00670). The exact baseline-composition term was −0.00106 (−0.00207 to −0.00018): six-month completers began with a slightly lower mean inflammatory score than the full baseline group. Thus the marginal contrast was more negative than the paired contrast, but the baseline-composition difference did not account for the observed decline.

**Figure 2:**
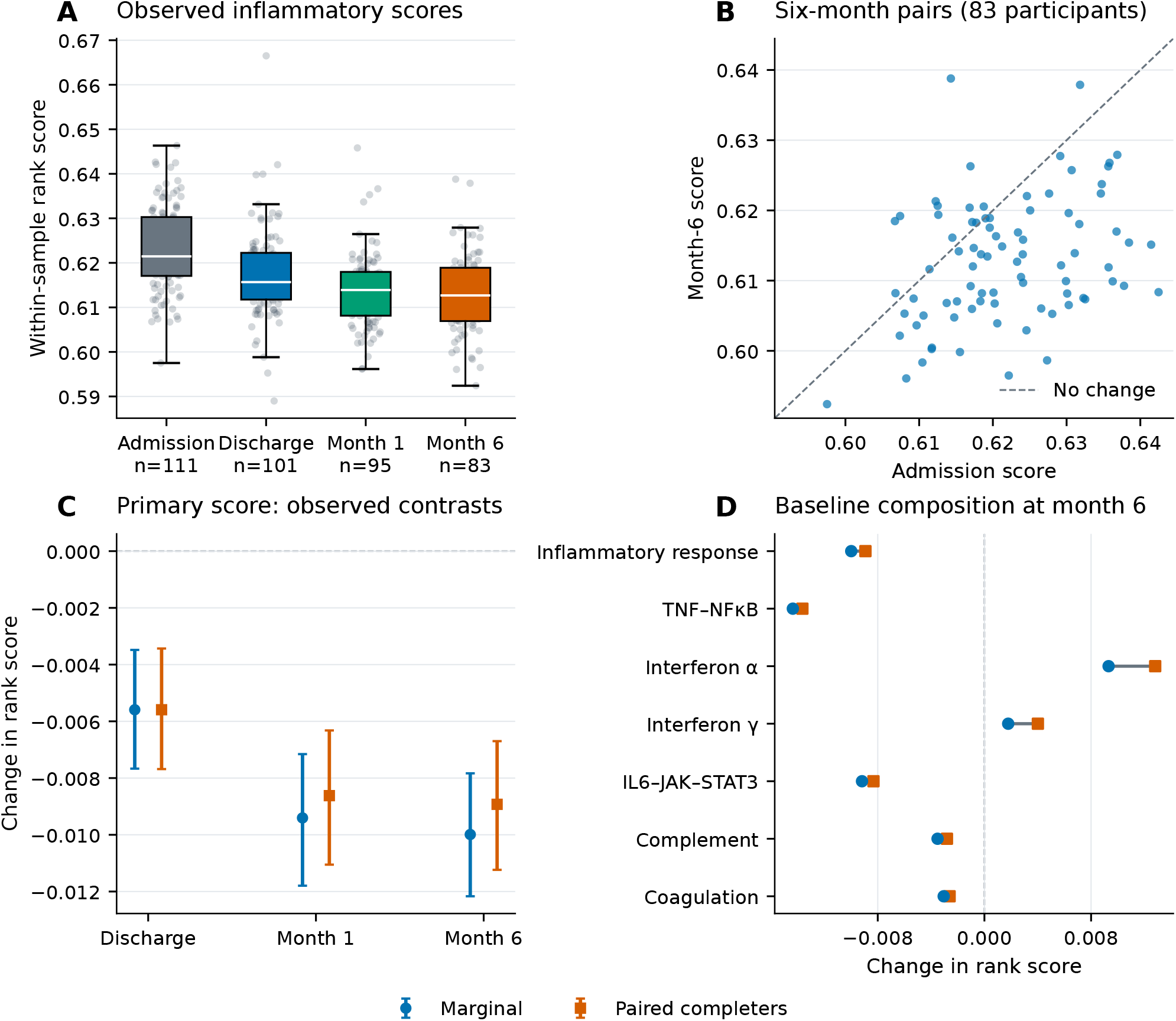
Observed transcriptomic change and the effect of baseline composition. (A) Inflammatory-response rank scores in the source cohort. Boxes show medians and interquartile ranges; whiskers extend to the most extreme observations within 1.5 interquartile ranges. Points show every assay, with horizontal jitter only. (B) Authenticated admission/month-6 pairs for 83 participants; the diagonal denotes equal scores. (C) Marginal change subtracts the mean of all 111 admission scores from the available follow-up mean. Paired-completer change subtracts each completer’s own admission score. Bars are 95% percentile intervals from 2,000 participant bootstrap draws. (D) The two contrasts for all seven fixed scores at month 6. Each connecting segment equals the completer admission mean minus the all-baseline mean; it is a baseline-composition component, not an uncertainty interval. Scores measure relative within-sample enrichment and are not clinical recovery outcomes. The shared legend applies to panels C and D.

The corresponding marginal contrasts were −0.00559 at discharge and −0.00941 at one month. At six months, TNF/NF-*κ*B and IL6/JAK/STAT3 scores also declined, whereas interferon-*α* and interferon-*γ* scores increased. These heterogeneous directions caution against describing all seven programs as a single immune-recovery axis. They are relative-expression summaries; changing cell composition or gene-background expression could contribute.

### 3.3 Baseline-score adjustment changes the primary estimate little

The nuisance-model diagnostics are summarized in Figure 4. Six-month availability discrimination was modest (held-out AUROC 0.602). The logistic Brier score was 0.1877 versus 0.1889 for the intercept baseline, while log loss was slightly worse (0.5723 versus 0.5654). Earlier-visit logistic AUROCs were 0.476 and 0.426, and their Brier scores were worse than the intercept baselines. Weak prediction neither demonstrates absence of selection nor establishes MAR. The permutation diagnostic placed the six-month AUROC within the null distribution’s upper tail rather than separating it decisively (eight of 99 permuted AUROCs were at least as large; corrected exceedance fraction 0.09).

Model-dependent changes, weights, and prespecified sensitivity settings are shown in Figure 5. At six months, the minimum held-out availability probability was 0.175; the largest observed inverse-probability weight was 5.70 and the effective sample size was 72.2, compared with 83 available assays. The primary inflammatory changes were −0.00971 for IPW, −0.00951 for g-computation, and −0.01004 for AIPW. Their 95% intervals were [−0.01225, −0.00738], [−0.01175, −0.00731], and [−0.01556, −0.00727], respectively. All 500 six-month nuisance refits succeeded; discharge had one failed refit out of 500 because its training data contained a degenerate availability outcome. That failure was retained in the audit log, and discharge intervals use 499 successful refits.

Across the eight prespecified settings, the primary AIPW point estimate ranged from −0.01054 to −0.00968. This is stability within the evaluated model family and measured baseline scores, not verification that missing follow-up outcomes are ignorable. In particular, the agreement of three MAR-based methods cannot identify outcomes under an unobserved-outcome-dependent selection process.

### 3.4 Broad nonidentification coexists with robustness within the prespecified SD range

Figure 3 separates the support interval from sampling uncertainty and standardized scenarios. The primary full-support completion interval was [−0.16460, 0.08765]. Its inclusion of zero means that score support alone does not identify the direction of the hypothetical full-baseline-cohort change. The interval’s width of 0.25225 equals the unavailable fraction by construction. This algebraic width is not a new biological result.

**Figure 3:**
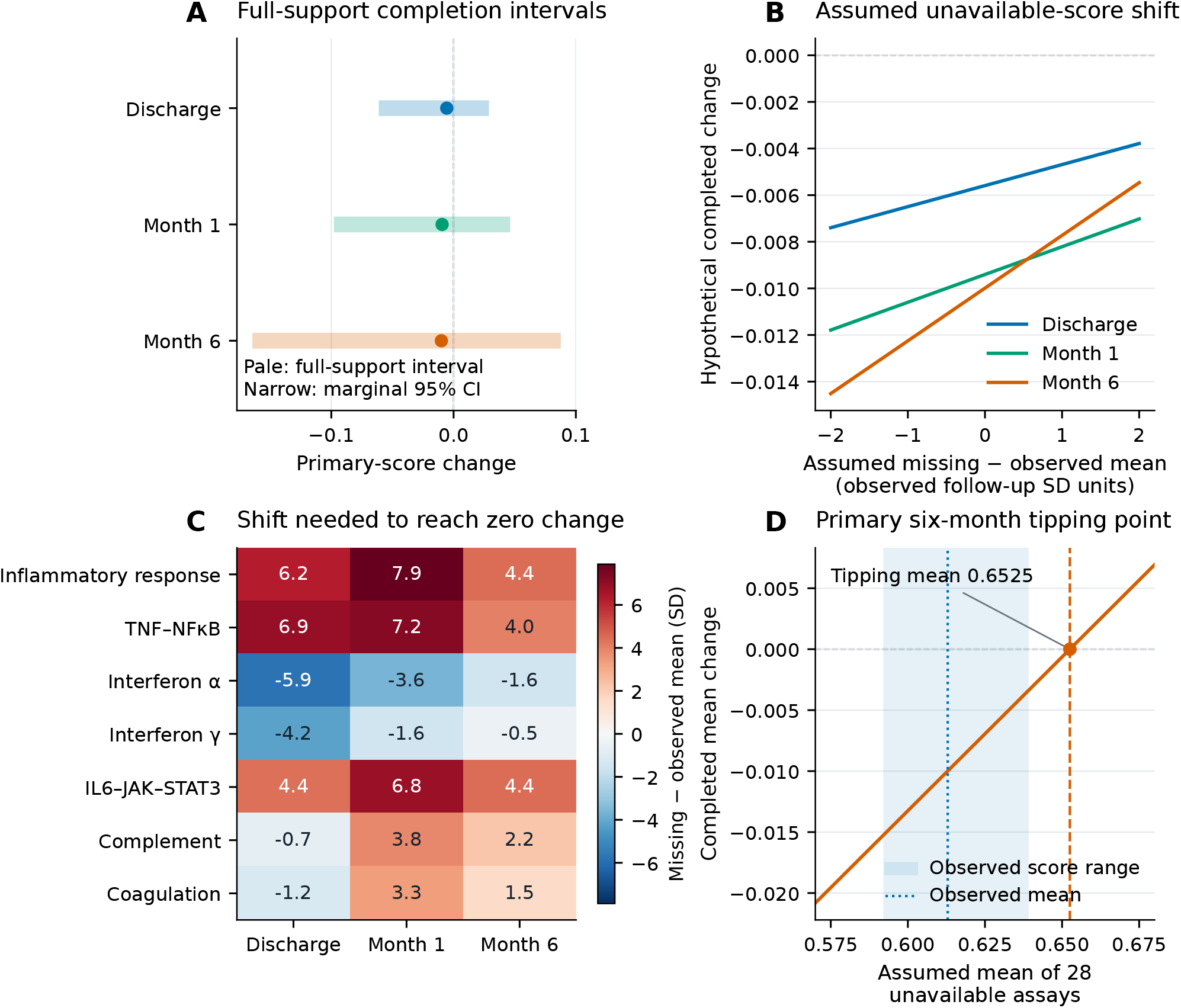
Identification intervals and explicit unavailable-assay assumptions. (A) Pale bands are full-support completion intervals for the primary inflammatory score, allowing the unavailable-assay mean anywhere in [0,1]. Narrow lines are observed marginal 95% bootstrap confidence intervals and points are marginal estimates. Identification intervals and sampling intervals answer different questions. (B) Completed-cohort change when the unavailable mean differs from the observed mean by minus two to plus two observed follow-up standard deviations (SD). These are assumption scenarios, not measured outcomes. The primary score remains negative throughout this grid. (C) Missing-minus-observed mean shift, in observed SD units, required to bring each marginal score change to zero. Secondary scores and visits are descriptive. (D) The primary month-6 tipping mean is 0.6525, versus an observed mean of 0.6129. The shaded observed score range is descriptive and is not imposed as a support bound. All completion results assume a defined assay outcome for every baseline participant; undefined outcomes after death are not resolved.

**Figure 4:**
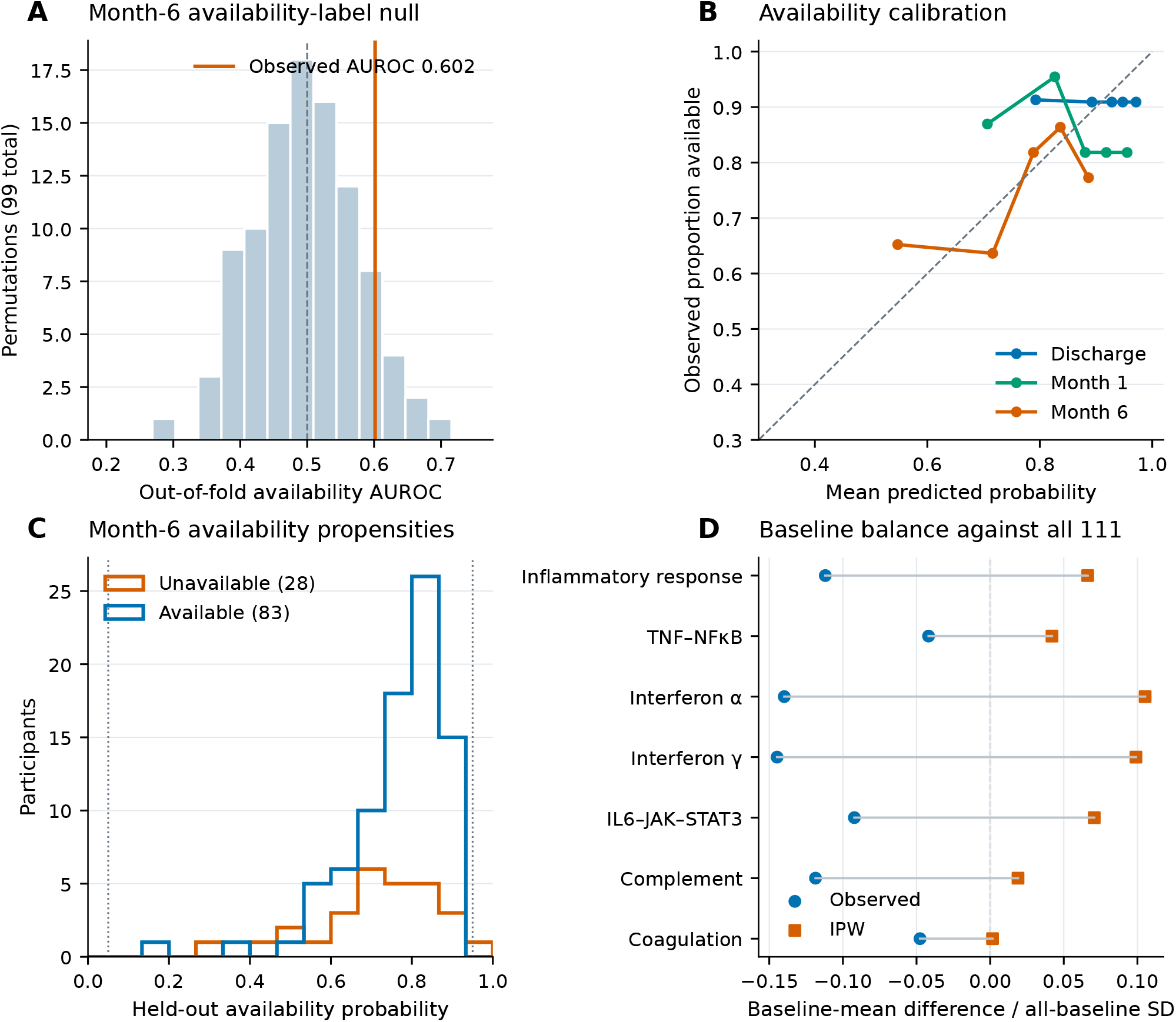
Held-out availability prediction and baseline-balance diagnostics. (A) Month-6 logistic availability AUROC compared with 99 patient-level availability-label permutations that preserve the observed count. The observed AUROC is 0.602; this diagnostic does not establish the natural missingness mechanism. (B) Held-out logistic predictions grouped into five approximately equal-sized bins separately at each visit. Points show bin mean predicted probability and observed availability; lines only connect bins. (C) Month-6 held-out propensity distributions in the 83 available and 28 unavailable participants. Dotted lines mark the primary clipping limits, 0.05 and 0.95. (D) Admission-score mean differences between available participants and all 111 baseline participants, before and after inverse-probability weighting, divided by the all-baseline SD. Connecting segments show the weighting-induced change in balance. Predictive performance and balance do not prove missing at random, and weighted balance need not improve for every score.

**Figure 5:**
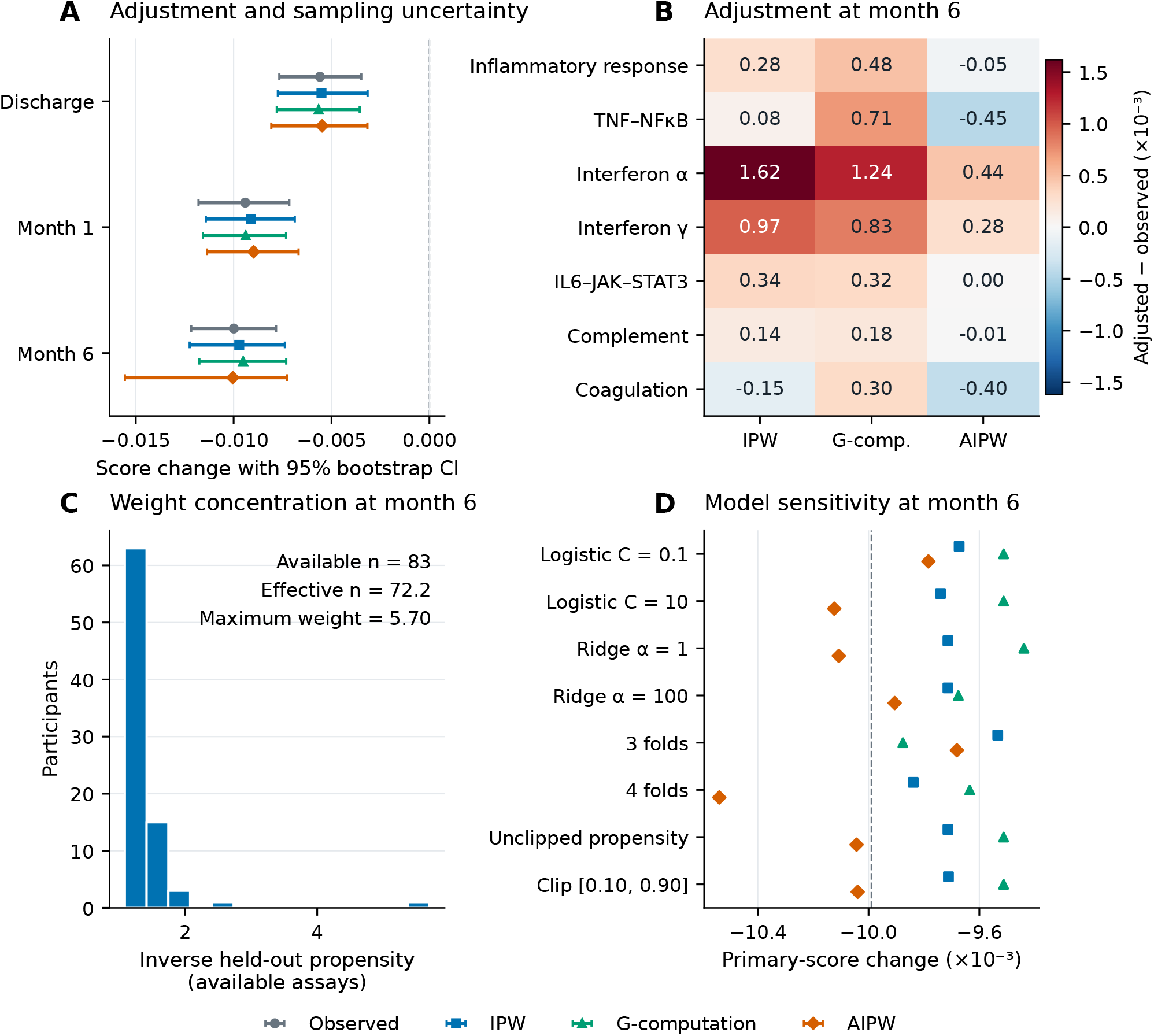
Assumption-dependent adjustment, precision and model sensitivity. (A) Primary-score changes from observed marginals, normalized inverse-probability weighting (IPW), g-computation and augmented inverse-probability weighting (AIPW). Bars are 95% percentile participant-bootstrap intervals. Observed estimates use 2,000 draws; nuisance-refit intervals use 500 draws, with 499 successful fits at discharge because one draw had degenerate training availability. (B) Month-6 adjustments minus the observed marginal estimate for all seven scores, multiplied by 1,000. (C) Distribution of inverse held-out propensities among the 83 month-6 available participants. The effective sample size is 72.2 and the maximum weight is 5.70. (D) Prespecified changes to logistic regularization, ridge regularization, fold count and propensity clipping. The vertical dashed line is the observed marginal estimate. These are sensitivity analyses, not externally selected models. Adjusted estimates require missing-at-random, positivity and model assumptions; agreement among estimators is not evidence that those assumptions hold. The shared legend applies to panels A and D.

More informative on the measured scale, a zero overall change required a mean score of 0.65253 among the 28 unavailable assays, compared with 0.61293 observed at six months. The difference of 0.03960 is 4.42 observed-follow-up SDs. The tipping mean exceeds the largest observed six-month score (0.63879), although observed extrema cannot restrict unmeasured assays. Over the prespecified *±*2-SD missing-mean range, the completed primary contrast remained negative, from −0.01451 to −0.00547. Consequently, these data do not support a claim that modest evaluated departures necessarily erase the inflammatory decline.

Robustness was program-specific. The six-month tipping shifts were 3.98 SD for TNF/NF-*κ*B, 4.42 SD for IL6/JAK/STAT3, and 2.19 SD for complement, versus −1.57 SD for interferon-*α, −*0.47 SD for interferon-*γ*, and 1.50 SD for coagulation. The latter three crossed zero somewhere in the *±*2-SD grid. These thresholds characterize assumptions required to change the sign; they neither estimate missing expression nor attach a probability to each scenario.

### 3.5 Artificial deletion exposes the limits of agreement between estimators

The artificial-deletion experiments (Figure 6) distinguish estimator bias from dispersion. Uniform deletion produced small average errors for the unweighted primary contrast. At the planned 25% deletion level, its RMSE was 0.00058, compared with 0.00055 for IPW and 0.00083 for AIPW. Under baseline-dependent deletion at that level, mean errors were −0.00099, 0.00045, and 0.00050, but the corresponding RMSEs were 0.00106, 0.00140, and 0.00155. Reduced average bias did not guarantee lower error across masks. With 50 masks per setting, these error summaries have limited Monte Carlo precision; no Monte Carlo uncertainty analysis was used to test differences between methods, and small RMSE differences are descriptive.

**Figure 6:**
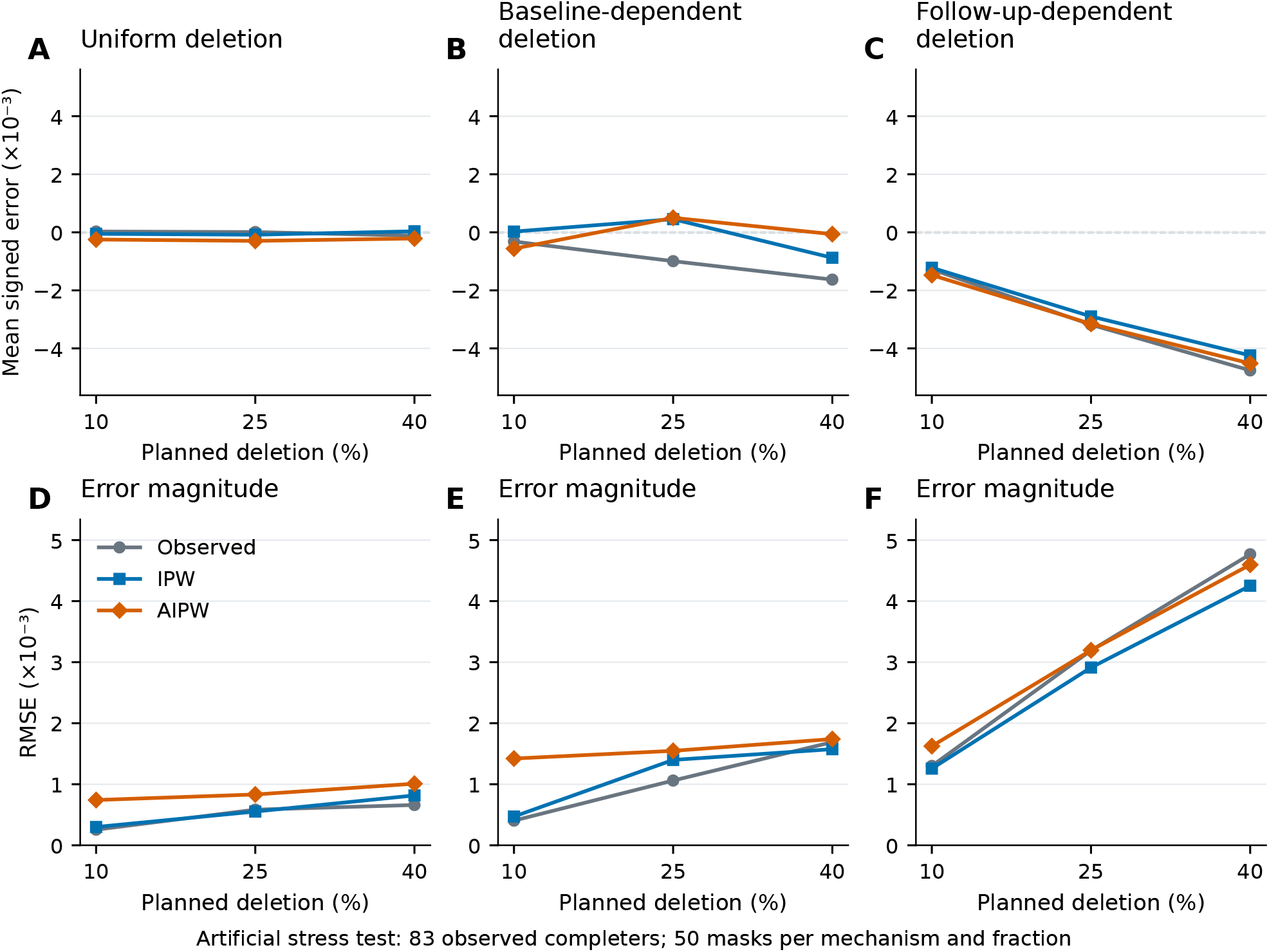
Artificial-deletion stress tests on the observed six-month parent set. The known-data parent consists of the 83 actual month-6 completers. Each deletion rule and planned fraction was repeated for 50 masks; these masks are not independent cohorts or new patients. Panels A-C show mean signed error and panels D-F show root mean squared error (RMSE) of the primary-score completed-cohort contrast relative to its known parent value. Columns represent uniform deletion, deletion depending on the baseline inflammatory score, and deletion depending on the follow-up inflammatory score. The planned 10%, 25% and 40% deletion fractions remove 8, 21 and 33 records, respectively. Observed-marginal, IPW and AIPW estimates are recomputed for each masked dataset. All panels within a row share a vertical scale. Follow-up-dependent deletion creates persistent error that baseline-only adjustment does not remove. These deliberately imposed mechanisms cannot diagnose why assays are absent in the original data.

When deletion depended on the follow-up inflammatory score, all three methods remained biased downward. At planned 25% deletion, mean errors were −0.00318 (unweighted), −0.00290 (IPW), and −0.00315 (AIPW). At planned 40% deletion, the corresponding errors were −0.00476, −0.00424, and −0.00452. Thus similar adjusted and unadjusted estimates can coexist with appreciable error under this specified mechanism. The experiment does not establish that actual unavailable participants resemble artificially removed completers, and its error scale cannot be transferred to the 28 genuinely absent assays.

### 3.6 Unlinked external data illustrate a narrower evidence claim

External marginal and conditional results are displayed in Figure 7. In the external whole-blood dataset, the inflammatory-response mean was 0.60479 at admission, 0.60231 at day 30, and 0.60020 at year one. These give marginal contrasts of −0.00248 and −0.00459. Because individual pairs are unavailable, these are comparisons of deposited assay groups, not average within-person changes. No confidence interval is labeled as a paired external validation.

**Figure 7:**
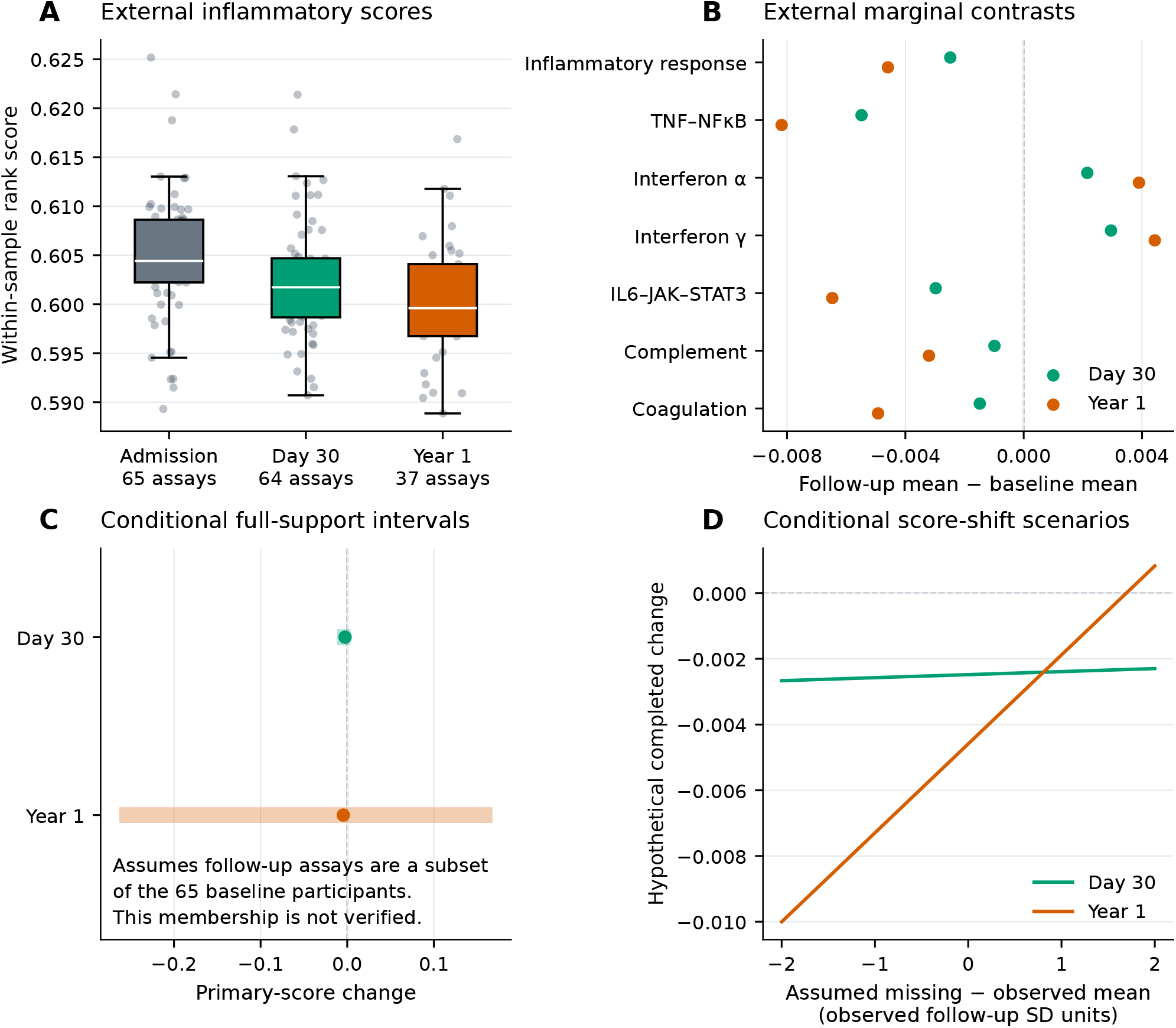
External marginal evidence and explicitly conditional completion scenarios. (A) Inflammatory-response score distributions for 65 admission, 64 day-30 and 37 year-1 external MI assays, after explicit technical-replicate removal. CAD comparator assays are excluded. Boxplot conventions match Figure 2. (B) External marginal contrasts for seven fixed scores, without paired inference or invented participant-level precision. (C) Conditional full-support intervals for the primary score; dots are the observed marginal follow-up-minus-baseline contrasts, not fitted missing-outcome estimates. These calculations assume that the 64 or 37 follow-up assays represent distinct members of the same 65-person baseline group. Public participant-level linkage and exact subset membership are not verified. (D) Corresponding conditional scenarios over a missing-minus-observed mean shift of minus two to plus two follow-up SD. Neither C nor D establishes external patient retention or validates the source propensity model. Different specimen types, platforms and follow-up horizons also limit direct cohort equivalence.

**Figure 8:**
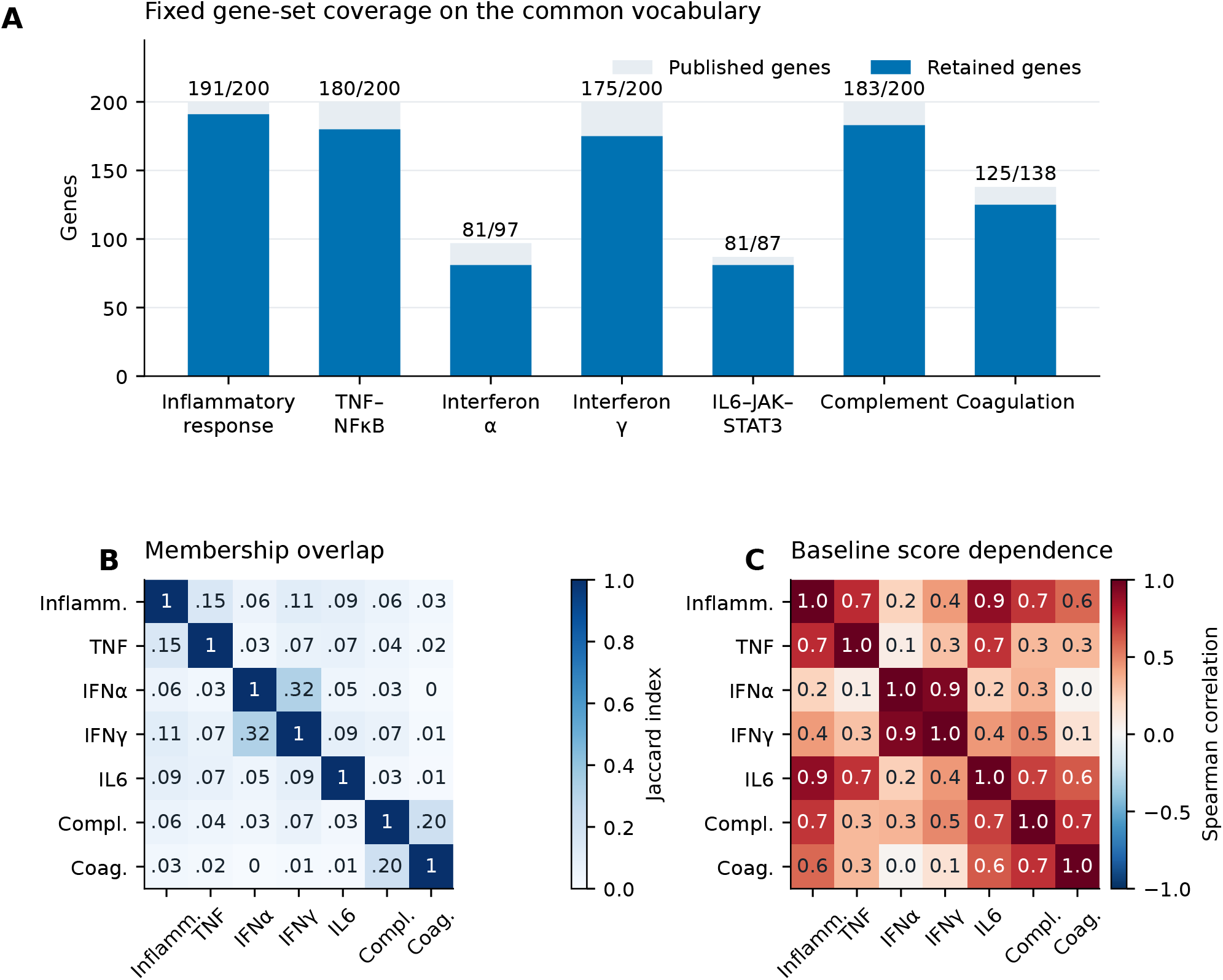
Measurement coverage and dependence of the fixed score panel. (A) Retained and published membership for the seven prespecified Hallmark sets, using the same 17,590-gene annotation-defined vocabulary in both cohorts. All sets retain at least 81 genes and at least 83.5% of published membership. (B) Jaccard overlap of retained gene memberships, defined as intersection divided by union. Compact cell annotations omit leading zeros; 1 denotes complete overlap. (C) Descriptive Spearman correlations between baseline source scores in 111 participants. This correlation display was added after the frozen analysis as an exploratory measurement diagnostic; it did not influence gene-set choice, fitting or primary inference. Correlated or overlapping scores are not independent biological replications. The fixed inflammatory-response score remains the primary endpoint; the other six scores provide prespecified descriptive sensitivity context.

If, additionally, all 37 year-one assays are assumed to be distinct members of the 65-person baseline array group, a zero completion contrast requires an unavailable-assay mean of 0.61085: 0.01065 above the observed mean, or 1.70 observed SDs. The corresponding conditional support interval is [−0.26314, 0.16763]. This conditional scenario is more sensitive within the evaluated SD scale than the source six-month primary result. Tissue composition, timing, clinical setting, and uncertain membership differ, so it cannot be interpreted as a failed replication of a six-month PBMC patient-level effect. Its central contribution is to show what missing linkage prevents the public data from resolving.

## 4 Discussion

The main finding is a distinction between empirical stability and assumption-free identification. In a linked public STEMI cohort, the inflammatory rank score declined among measured follow-up samples, the baseline-composition contribution was small relative to the observed change, and several prespecified MAR estimators gave similar results. The primary direction also persisted when the unavailable-assay mean was moved by up to two observed SDs. Nevertheless, unrestricted completion over the score support included both signs. Reporting only the broad interval would understate the observed robustness; reporting only the narrow bootstrap interval would overstate what is known about the complete baseline cohort.

The study’s contribution is an empirical audit of an established evidence source, not a new missing-data theory. Prior publications have identified temporal immune programs, candidate heart-failure markers, and recovery-associated signatures in these and related cohorts[1–4]. Our fixed Hallmark measurement avoids optimizing a recovery signature on these follow-up outcomes. Relative to the recovery-signature task of Cheng et al. [4], our analysis changes the estimand to hypothetical completion of the authenticated baseline cohort: how much information about unavailable assays is required to change that population-level conclusion? Because the published recovery signature was not reimplemented, these findings neither reproduce nor refute its performance. This distinction also separates the present work from classification benchmarks, where discrimination between case and comparison groups is the target[10]. A molecular difference may be reproducible among observed specimens without identifying a clinical recovery state for every enrolled individual.

Clinical endpoint studies clarify why this distinction matters. A nationwide observational study of intra-aortic balloon pump use in patients with MI and Killip class III or IV undergoing PCI evaluated in-hospital mortality[18]. That clinical endpoint and treatment comparison differ from the relative blood-expression contrasts examined here. The registry findings neither validate our transcriptomic measure nor identify the survival status of participants without later assays. Conversely, our score decline cannot be used to infer a mortality benefit or the effectiveness of circulatory support.

The proposed tri-ecological vascular-device Suitcordance framework distinguishes mechanical, cellular, and physicochemical-immune aspects of the device–vessel environment[19]. Our relative blood-expression measurements sample a limited molecular readout and cannot establish coordinated recovery across those domains. This conceptual distinction motivates pairing future molecular trajectories with independently measured vascular-tissue and clinical outcomes.

The source primary result was more robust than several secondary programs. Interferon scores and coagulation could change direction under smaller standardized shifts, despite being measured in the same participants. Missing fractions alone therefore do not rank the robustness of all biological summaries. A useful report needs the observed effect, missing fraction, score variation, and tipping threshold together. Conversely, a threshold expressed in observed SDs is not a calibrated measure of biological plausibility: the observed group may have a truncated distribution, and program scores have different composition and measurement properties. The *±*2-SD range was an analysis convention, not a validated clinical bound.

Observable selection adjustment was limited both by information and by sample size. Seven baseline rank scores may miss determinants associated with clinical status, treatment, technical quality, access to follow-up, or survival. Availability prediction was weak, and weighting reduced the effective sample size. More elaborate machine learning would not itself identify the missing-outcome mechanism. In artificial deletion, MAR estimators could reduce average error for a baseline-dependent mechanism while increasing dispersion; none removed the bias caused by the specified follow-up-dependent mechanism.

This demonstrates a failure mode of these implementations on this finite parent, rather than a universal ranking of estimators. The disclosed difference between the planned rank-weighting wording and the executed standardized-score weights further limits these experiments to the implemented stress tests; it does not affect the observed source-cohort estimates. With only 50 masks per setting and a selected completer parent, broader claims would require additional simulation designs and independently linked cohorts.

The external dataset exposes a practical limitation of public longitudinal omics: visit labels without participant linkage are insufficient for paired inference. Technical-replicate removal does not recover a missing roster, and the number of later arrays does not prove that they are a subset of the baseline array group. Conditional scenarios are useful for displaying the consequences of a stated assumption, but should not acquire the language of verified retention or independent MAR validation. Depositing pseudonymous longitudinal identifiers, assay-selection rules, scheduled and measured visits, and reasons for nonavailability would be more valuable for this question than adding another retrospective model to the same incomplete record.

Several limitations constrain interpretation. First, only one cohort supports authenticated participant-level analysis. Second, scores depend on official processed microarrays and annotation decisions; all-array normalization and cross-platform measurement differences are inherited. Third, relative rank enrichment cannot separate cell-composition shifts from intracellular regulation, and the common vocabulary changes what a gene-set score means. Fourth, the completion target assumes an assay is defined for every participant; unknown survival and absence reasons prevent a clinically interpretable survivor or mortality-adjusted estimand[15]. Fifth, broad support bounds are intentionally uninformative when small expression changes coexist with appreciable absence; they are not evidence of actual reversal. Sixth, model-based intervals condition on the score definition and evaluated nuisance models. Finally, the public data and their earlier biological findings were known before the internal protocol, and the audit is retrospective. None of these analyses validates a biomarker for treatment selection, establishes intervention efficacy, or demonstrates improvement in an autonomous scientific agent.

For future vascular transcriptomic studies, the actionable conclusion is to publish the target population alongside a recovery estimate. Paired change, marginal change, baseline composition, model-dependent completion, and missing-mean tipping thresholds answer different questions and can be reported together. In the evaluated source cohort, this joint report supports a measured inflammatory decline that is stable throughout the prespecified *±*2-observed-SD missing-mean scenarios, while preserving uncertainty about unavailable assays and about broader clinical recovery.

## Supporting information

Supplementary methods and results

Manuscript LaTeX source files

Reproducibility code and derived results

## Data and code availability

Official processed data are available from GEO accessions GSE59867 and GSE123342, with platform annotations GPL6244 and GPL17586. The accompanying reproducibility archive includes acquisition scripts, source URLs and hashes, sample inventories, fixed gene memberships, the timestamped internal analysis specification and clarification, analysis and validation code, saved numerical outputs, and figure-generation code. Original third-party data remain subject to their source terms. A separate public archival identifier has not yet been assigned to this study’s code package.

## Ethics, consent, and registration

This analysis uses existing public expression records without recruitment, intervention, or access to direct personal identifiers. Ethics and consent for the original cohorts are reported in their source publications. The submitting author confirms that applicable requirements for this public-data secondary analysis have been addressed. No new institutional ethics approval or exemption is claimed. This is not a clinical trial, and no trial registration is claimed. The internal analysis specification was not prospectively registered.

## Author contributions

Lingsen You, Zisu Peng, and Yujun Guo contributed equally to this work. All authors reviewed and approved the manuscript and agreed to its submission.

## DECLARATION OF GENERATIVE AI AND AI-ASSISTED TECHNOLOGIES

Codex assisted with LaTeX assembly, prose editing, and consistency checks. The authors reviewed and revised all AI-assisted text and take responsibility for the submitted manuscript.

