## Supplementary methods and results for "Follow-up availability and the robustness of immune transcriptomic recovery after myocardial infarction"

These materials accompany *Follow-up availability and the robustness of immune transcriptomic recovery after myocardial infarction.* All estimates are computed from the saved public-data analysis. Hypothetical completion scenarios and artificial deletion are labeled separately from observed measurements. Rounded entries are for display; machine-readable CSV files retain full precision.

#### S1. Dataset inventory and units of analysis

The linked source consists of 111 STEMI participants in GSE59867. Its 390 STEMI arrays are repeated measurements, not 390 independent people. All 436 retained arrays were scored, but the 46 stable-CAD controls are excluded from the longitudinal estimand. The second deposit has 192 arrays; removal of four explicitly identified technical replicates leaves 188 scored arrays. Its recovery summaries use 166 MI assays, excluding 22 CAD controls. The 65 admission MI assays are not combined with later assays to form a count of independent participants. The external parent study included mixed acute MI; the exact STEMI/NSTEMI composition of the deposited microarray subset is not established here. The two studies are never pooled into an asserted unique-patient total.

| Deposit / specimen | Deposited | Scored arrays | Recovery-analysis role |
|---|---|---|---|
| GSE59867 / PBMC | 436 | 436 | 390 STEMI assays; 111 linked participants |
| GSE123342 / whole blood | 192 | 188 | 166 MI assays; unlinked visits |

**Platforms and measurement.** GSE59867 uses Affymetrix Human Gene 1.0 ST (GPL6244); GSE123342 uses Human Transcriptome Array 2.0 (GPL17586). Official processed matrices are used without redoing raw-array normalization. The endpoint is a relative transcriptomic rank score, not myocardial function, mortality, clinical recovery, or treatment benefit.

**Follow-up and censoring.** Source titles authenticate person/visit links. The data do not provide an individual reason for each absent assay. No censoring time, complete death roster, or longitudinal treatment exposure is available for analysis. External hybridization identifiers identify assays; adjacency and expression similarity were not used to invent patient pairs.

**Original ethics information.** The source publications report original ethics approval and informed consent (PMCID PMC4432772; PMC6922070). This describes the historical studies; it is not a new approval, exemption, or institutional determination for the current secondary analysis.

**Reuse conditions.** NCBI encourages molecular-data reuse and imposes no restrictions itself, while submitters may retain rights. Hallmark sets were retrieved under the published MSigDB CC BY 4.0 terms. The manuscript license does not override third-party terms. Official policy records, retrieval dates, file hashes, and a complete structured inventory are in `evidence/data_registry.json` and `dataset_use_conditions.json`.

#### S2. Follow-up structure and duplicate handling

| Admission/discharge/M1/M6 pattern | Participants |
|---|---|
| 1000 | 3 |
| 1001 | 2 |
| 1011 | 5 |
| 1100 | 8 |
| 1101 | 3 |
| 1110 | 17 |
| 1111 | 73 |

A digit of 1 denotes an available deposited assay; 0 denotes absence. There are ten participants with an unavailable interim assay followed by a later available assay. Visit-specific availability models therefore do not assume monotone dropout. The source denominator is always 111 at each visit.

| Visit | GSE59867 STEMI assays | GSE123342 MI assays |
|---|---|---|
| Admission | 111 | 65 |
| Discharge | 101 | Not sampled |
| One month / day 30 | 95 | 64 |
| Six months | 83 | Not sampled |
| One year | Not sampled | 37 |

##### Explicitly excluded external technical replicates

| Replicate GSM | Replicate hybridization | Original GSM | Original hybridization |
|---|---|---|---|
| GSM3501245 | hyb18756 | GSM3501376 | hyb20838 |
| GSM3501246 | hyb18757 | GSM3501377 | hyb20839 |
| GSM3501265 | hyb18778 | GSM3501320 | hyb20782 |
| GSM3501266 | hyb18779 | GSM3501321 | hyb20783 |

The original external report describes a larger 143-person discovery cohort and a 65-person microarray branch. Parent-cohort death or loss counts cannot be assigned to the microarray subset. Consequently, 64/65 and 37/65 are array-count ratios, not authenticated participant-retention estimates. The hypothetical completion calculations explicitly assume later assays are distinct members of the 65-person baseline group; that subset assumption remains unverified. Stable-CAD assays are contextual controls and do not supply missing MI follow-ups.

#### S3. Fixed score construction, coverage, and overlap

Probes with exactly one valid symbol were retained, and the median was taken across probes mapping to the same symbol within each array. The annotation-and-assay-presence intersection contains 17,590 symbols. This vocabulary does not depend on effect sizes or model performance. Average-tie ranks are calculated within each sample. Seven Hallmark memberships were fixed before scoring. No set failed the specified coverage threshold of at least 20 genes and at least half of its original membership.

| Program | Original genes | Retained genes | Coverage |
|---|---:|---:|---:|
| Inflammatory | 200 | 191 | 95.5% |
| TNF/NF-kB | 200 | 180 | 90.0% |
| IFN-alpha | 97 | 81 | 83.5% |
| IFN-gamma | 200 | 175 | 87.5% |
| IL6/JAK/STAT3 | 87 | 81 | 93.1% |
| Complement | 200 | 183 | 91.5% |
| Coagulation | 138 | 125 | 90.6% |

Coverage is identical for both datasets because the same common vocabulary is used. For $G$ genes and $k$ members, the score is $(\sum_{g \in A} \text{rank}(g) - k(k+1)/2)/[k(G-k)]$. It equals the fraction of member/background comparisons in which the member is more highly expressed, with ties contributing one half. It is bounded by zero and one. Positive shifts can arise from member or background changes and altered cell mixture; the score is not a cell-intrinsic pathway activation measurement.

##### Retained gene overlap (counts)

| Program | 1 | 2 | 3 | 4 | 5 | 6 | 7 |
|---|---:|---:|---:|---:|---:|---:|---:|
| 1 Inflammatory | 191 | 49 | 15 | 36 | 22 | 21 | 9 |
| 2 TNF/NF-kB | 49 | 180 | 7 | 23 | 18 | 15 | 6 |
| 3 IFN-alpha | 15 | 7 | 81 | 62 | 7 | 8 | 1 |
| 4 IFN-gamma | 36 | 23 | 62 | 175 | 21 | 22 | 4 |
| 5 IL6/JAK/STAT3 | 22 | 18 | 7 | 21 | 81 | 7 | 2 |
| 6 Complement | 21 | 15 | 8 | 22 | 7 | 183 | 52 |
| 7 Coagulation | 9 | 6 | 1 | 4 | 2 | 52 | 125 |

Program numbering follows the coverage table. The diagonal gives retained size; overlap means these seven summaries are correlated and must not be counted as seven independent replications. Full retained memberships and Jaccard overlaps are saved in `results/retained_gene_sets.json` and `gene_set_overlap.csv`.

#### S4. Provenance, checksums, and acquisition conditions

All input downloads below occurred on 27 October 2026 (UTC). SHA-256 identifies the exact analyzed bytes, not the current content of a changing URL. Annotation hashes identify the first complete platform table extracted from the official family stream, not the entire family download. Individual URLs and timestamps are retained in the acquisition JSON files.

| Resource | SHA-256 (two contiguous halves) |
|---|---|
| GSE59867 matrix | `0f90638ce4e8b2e33b4ee2cf8c364164`<br>`ca3e538155c9b2e8e56cd0ebc69986a5` |
| GSE123342 matrix | `5ac7e823f09e1a3873c8e73094bc246b`<br>`b0300f2add5e7ec1f04680fd6d4b6815` |
| GPL6244 annotation | `fec006e4ce50b9652e57dfd1a9662dd2`<br>`d3d8d52dd4080ddd7c588a1cfebb2ca6` |
| GPL17586 annotation | `41aef177aebe656c82cdff02ae070372`<br>`a60f4ba3b3a96d933a8a4de9bb4ff39d` |
| Inflammatory membership | `35203e49f5dbc3d0ada06ef005bc555a`<br>`07666b860a05ac0de12bc6a6c19773ae` |
| TNF/NF-kB membership | `ade1ab9b67e047ceb70f8b8a06c65c66`<br>`31e17f06b10c1b0283f962be848d2954` |
| IFN-alpha membership | `2a3d93f3170bc9866ced1135bdde9870`<br>`f18294f7ad3ae6cb3732dc3680483455` |
| IFN-gamma membership | `845aabdb9ff04fb62bbb1bc8a527654c`<br>`549075852c84f31d64f50d9e43134353` |
| IL6/JAK/STAT3 membership | `449e66144deb5f09187afe823b20e94a`<br>`1e607a86c29707603e2088b7fa5e7d83` |
| Complement membership | `b107622fa1384c10a15849470a5cac37`<br>`44cbb0672c6af7df3f748817a217f12f` |
| Coagulation membership | `9f4497576123ed73ed430335a24c4d91`<br>`ec9a6a8622a3f68dc73c67238fd3d335` |
| Frozen protocol | `74c50c831d46b0e7e0fe918cd95e870c`<br>`a17474c104ebf0a793fcd94a2e27f02a` |
| Score table | `77bf8eb915b0b4f61689b088e0754dc6`<br>`b54f36c2c8ad765b1e8ac83cd0051d88` |

Official series: `https://www.ncbi.nlm.nih.gov/geo/query/acc.cgi?acc=GSE59867`; `https://www.ncbi.nlm.nih.gov/geo/query/acc.cgi?acc=GSE123342`.

Data-use policy: `https://www.ncbi.nlm.nih.gov/home/about/policies/`. Gene-set terms: `https://www.gsea-msigdb.org/gsea/msigdb_license_terms.jsp`. All seven selected sets are Hallmark members, not KEGG or BioCarta sets with collection-specific restrictions. GSE28454 and GSE29111 were downloaded only during feasibility assessment; neither contributes observations or results to this paper.

#### S5. Estimands, decomposition, and identification

Let $X_i$ be the admission score, $Y_i$ the follow-up score when defined, and $R_i$ indicate an available follow-up assay. Write $r = n^{-1}\sum_i R_i$, $\bar{X} = n^{-1}\sum_i X_i$, and $\bar{Y}_R = \sum_i R_i Y_i / \sum_i R_i$. The observed marginal contrast and paired-completer contrast differ:

$$D = \bar{Y}_R - \bar{X},$$
$$C = \frac{\sum_i R_i(Y_i - X_i)}{\sum_i R_i},$$
$$D - C = \frac{\sum_i R_i X_i}{\sum_i R_i} - \bar{X}.$$

The last term is an exact baseline-composition decomposition, not a causal estimate of selection bias. It does not reveal unmeasured future scores.

If every baseline participant is assumed to have a defined hypothetical follow-up score and $u$ denotes the mean among unavailable assays, the completed contrast is

$$\Delta(u) = r\bar{Y}_R + (1 - r)u - \bar{X}.$$

For $0 \le u \le 1$, its endpoints are $L = r\bar{Y}_R - \bar{X}$ and $U = L + 1 - r$. Thus $U - L = 1 - r$ identically. The completion interval is not a confidence interval. Crossing zero establishes only that the sign is not identified by the observed values and the full support restriction. It is neither proof that unavailable participants differ nor evidence that recovery is absent. Zero is attained at

$$u^* = \frac{\bar{X} - r\bar{Y}_R}{1 - r}, \qquad \delta^* = u^* - \bar{Y}_R = -\frac{D}{1 - r}.$$

Values of $u^*$ outside $[0, 1]$ are unattainable. Standardizing $\delta^*$ by the observed follow-up SD describes an assumption strength, not an observed missing outcome. Prespecified scenarios use $u = \text{clip}(\bar{Y}_R + d\, s_R, 0, 1)$ for $d \in [-2, 2]$.

##### MAR estimators and uncertainty

The propensity $\widehat{p}_i$ and outcome regression $\widehat{m}_i$ are obtained out of fold. The primary propensity is clipped to $[0.05, 0.95]$; outcome predictions are clipped to $[0, 1]$. Hajek IPW normalizes observed inverse-probability weights. G-computation averages all predictions. AIPW averages $\widehat{m}_i + R_i(Y_i - \widehat{m}_i)/\widehat{p}_i$ and subtracts $\bar{X}$. AIPW itself is not truncated. These procedures require the stated MAR, positivity and model conditions; they do not diagnose the missingness mechanism.

Observed-estimate intervals use 2,000 participant bootstrap draws. Adjusted intervals use 500 draws with nuisance refitting; repeated draws of the same original participant retain the same fold. Discharge replicate 238 has degenerate training availability and is recorded as a failure, leaving 499 adjusted draws; M1 and M6 each retain 500. Percentile intervals condition on the processed arrays and fixed score definitions.

##### Scores after death may be undefined

A future blood score is not defined in the same way after death. Without individual survival information, hypothetical completion does not identify a survivor-average or mortality estimand. No numerical missing-data procedure resolves this limitation. These calculations must not be interpreted as effects on survival, heart failure, or a clinical intervention.

#### S6. Complete observed source estimates

Each entry is a point estimate followed by a 95% participant-bootstrap percentile interval (2,000 draws). All values are changes in the bounded score, not percentages. The primary endpoint is inflammatory response at M6. The remaining visits/programs are descriptive secondary analyses. No multiplicity-adjusted significance claims are made.

| Visit / program | Marginal contrast $D$ | Paired contrast $C$ | Baseline composition |
|---|---:|---:|---:|
| Discharge / Inflammatory | −0.0056 [−0.0077, −0.0035] | −0.0056 [−0.0077, −0.0034] | −0.0000 [−0.0005, 0.0004] |
| Discharge / TNF/NF-kB | −0.0105 [−0.0145, −0.0062] | −0.0105 [−0.0146, −0.0061] | 0.0001 [−0.0007, 0.0008] |
| Discharge / IFN-alpha | 0.0159 [0.0103, 0.0216] | 0.0171 [0.0117, 0.0227] | −0.0012 [−0.0039, 0.0008] |
| Discharge / IFN-gamma | 0.0074 [0.0035, 0.0114] | 0.0082 [0.0045, 0.0122] | −0.0008 [−0.0023, 0.0003] |
| Discharge / IL6/JAK/STAT3 | −0.0040 [−0.0061, −0.0018] | −0.0040 [−0.0060, −0.0018] | −0.0000 [−0.0004, 0.0004] |
| Discharge / Complement | 0.0004 [−0.0007, 0.0014] | 0.0004 [−0.0008, 0.0015] | −0.0000 [−0.0002, 0.0002] |
| Discharge / Coagulation | 0.0009 [−0.0008, 0.0026] | 0.0005 [−0.0011, 0.0022] | 0.0004 [−0.0000, 0.0009] |
| M1 / Inflammatory | −0.0094 [−0.0118, −0.0072] | −0.0086 [−0.0111, −0.0063] | −0.0008 [−0.0017, −0.0000] |
| M1 / TNF/NF-kB | −0.0144 [−0.0185, −0.0102] | −0.0130 [−0.0173, −0.0084] | −0.0014 [−0.0031, −0.0001] |
| M1 / IFN-alpha | 0.0121 [0.0067, 0.0176] | 0.0114 [0.0060, 0.0169] | 0.0007 [−0.0014, 0.0029] |
| M1 / IFN-gamma | 0.0037 [0.0000, 0.0077] | 0.0035 [−0.0003, 0.0075] | 0.0002 [−0.0011, 0.0014] |
| M1 / IL6/JAK/STAT3 | −0.0084 [−0.0109, −0.0060] | −0.0078 [−0.0104, −0.0053] | −0.0006 [−0.0013, −0.0000] |
| M1 / Complement | −0.0031 [−0.0043, −0.0020] | −0.0030 [−0.0042, −0.0018] | −0.0002 [−0.0006, 0.0003] |
| M1 / Coagulation | −0.0039 [−0.0056, −0.0022] | −0.0034 [−0.0051, −0.0016] | −0.0005 [−0.0012, 0.0001] |
| M6 / Inflammatory | −0.0100 [−0.0122, −0.0078] | −0.0089 [−0.0112, −0.0067] | −0.0011 [−0.0021, −0.0002] |
| M6 / TNF/NF-kB | −0.0144 [−0.0182, −0.0106] | −0.0137 [−0.0177, −0.0097] | −0.0007 [−0.0027, 0.0011] |
| M6 / IFN-alpha | 0.0093 [0.0035, 0.0149] | 0.0128 [0.0074, 0.0182] | −0.0035 [−0.0071, −0.0002] |
| M6 / IFN-gamma | 0.0018 [−0.0019, 0.0055] | 0.0040 [0.0004, 0.0075] | −0.0022 [−0.0044, −0.0001] |
| M6 / IL6/JAK/STAT3 | −0.0092 [−0.0112, −0.0073] | −0.0083 [−0.0104, −0.0064] | −0.0009 [−0.0018, 0.0000] |
| M6 / Complement | −0.0035 [−0.0048, −0.0022] | −0.0028 [−0.0042, −0.0015] | −0.0007 [−0.0013, −0.0001] |
| M6 / Coagulation | −0.0031 [−0.0048, −0.0012] | −0.0026 [−0.0045, −0.0008] | −0.0004 [−0.0015, 0.0007] |

The interval for each component was computed directly on the same participant-resampling scheme. Subtracting two marginal interval endpoints does not reconstruct the interval for a difference. These three columns answer different questions even when their numerical values are close. Full precision and the six available bootstrap measures are supplied in `source_estimates.csv` and `bootstrap_intervals.csv`.

#### S7. Complete MAR-adjusted source estimates

Point estimates are followed by 95% refitted participant-bootstrap percentile intervals. IPW denotes normalized inverse-probability weighting; G-comp denotes outcome-regression standardization; AIPW denotes augmented inverse-probability weighting. Every contrast subtracts the entire admission mean. These are assumption-dependent estimates, not observed recovery measurements in unavailable participants.

| Visit / program | IPW | G-comp | AIPW |
|---|---|---|---|
| Discharge / Inflammatory | $-0.0055$ $[-0.0077, -0.0032]$ | $-0.0057$ $[-0.0078, -0.0036]$ | $-0.0055$ $[-0.0081, -0.0032]$ |
| Discharge / TNF/NF-kB | $-0.0105$ $[-0.0150, -0.0063]$ | $-0.0106$ $[-0.0148, -0.0068]$ | $-0.0103$ $[-0.0153, -0.0061]$ |
| Discharge / IFN-alpha | $0.0178$ $[0.0117, 0.0319]$ | $0.0162$ $[0.0107, 0.0216]$ | $0.0176$ $[0.0118, 0.0291]$ |
| Discharge / IFN-gamma | $0.0085$ $[0.0041, 0.0161]$ | $0.0076$ $[0.0037, 0.0113]$ | $0.0085$ $[0.0042, 0.0148]$ |
| Discharge / IL6/JAK/STAT3 | $-0.0041$ $[-0.0068, -0.0018]$ | $-0.0041$ $[-0.0063, -0.0019]$ | $-0.0040$ $[-0.0069, -0.0017]$ |
| Discharge / Complement | $0.0004$ $[-0.0007, 0.0018]$ | $0.0003$ $[-0.0008, 0.0014]$ | $0.0004$ $[-0.0009, 0.0015]$ |
| Discharge / Coagulation | $0.0005$ $[-0.0022, 0.0019]$ | $0.0008$ $[-0.0006, 0.0023]$ | $0.0004$ $[-0.0041, 0.0020]$ |
| M1 / Inflammatory | $-0.0091$ $[-0.0114, -0.0069]$ | $-0.0094$ $[-0.0116, -0.0073]$ | $-0.0090$ $[-0.0114, -0.0067]$ |
| M1 / TNF/NF-kB | $-0.0140$ $[-0.0181, -0.0099]$ | $-0.0145$ $[-0.0184, -0.0107]$ | $-0.0138$ $[-0.0181, -0.0095]$ |
| M1 / IFN-alpha | $0.0126$ $[0.0068, 0.0192]$ | $0.0117$ $[0.0064, 0.0172]$ | $0.0130$ $[0.0066, 0.0203]$ |
| M1 / IFN-gamma | $0.0042$ $[-0.0002, 0.0086]$ | $0.0035$ $[-0.0000, 0.0072]$ | $0.0045$ $[-0.0002, 0.0095]$ |
| M1 / IL6/JAK/STAT3 | $-0.0081$ $[-0.0107, -0.0059]$ | $-0.0084$ $[-0.0108, -0.0064]$ | $-0.0080$ $[-0.0107, -0.0057]$ |
| M1 / Complement | $-0.0029$ $[-0.0041, -0.0016]$ | $-0.0031$ $[-0.0041, -0.0019]$ | $-0.0028$ $[-0.0041, -0.0015]$ |
| M1 / Coagulation | $-0.0035$ $[-0.0052, -0.0017]$ | $-0.0037$ $[-0.0053, -0.0019]$ | $-0.0034$ $[-0.0051, -0.0015]$ |
| M6 / Inflammatory | $-0.0097$ $[-0.0122, -0.0074]$ | $-0.0095$ $[-0.0118, -0.0073]$ | $-0.0100$ $[-0.0156, -0.0073]$ |
| M6 / TNF/NF-kB | $-0.0143$ $[-0.0189, -0.0100]$ | $-0.0137$ $[-0.0172, -0.0098]$ | $-0.0148$ $[-0.0244, -0.0100]$ |
| M6 / IFN-alpha | $0.0109$ $[0.0044, 0.0173]$ | $0.0106$ $[0.0045, 0.0159]$ | $0.0098$ $[-0.0054, 0.0168]$ |
| M6 / IFN-gamma | $0.0028$ $[-0.0016, 0.0068]$ | $0.0026$ $[-0.0012, 0.0061]$ | $0.0021$ $[-0.0073, 0.0068]$ |
| M6 / IL6/JAK/STAT3 | $-0.0088$ $[-0.0108, -0.0067]$ | $-0.0089$ $[-0.0109, -0.0069]$ | $-0.0092$ $[-0.0133, -0.0068]$ |
| M6 / Complement | $-0.0034$ $[-0.0050, -0.0018]$ | $-0.0033$ $[-0.0047, -0.0020]$ | $-0.0035$ $[-0.0067, -0.0019]$ |
| M6 / Coagulation | $-0.0032$ $[-0.0060, -0.0010]$ | $-0.0027$ $[-0.0044, -0.0009]$ | $-0.0035$ $[-0.0089, -0.0012]$ |

Propensity predictors are the seven fixed admission scores, standardized using training folds only; logistic $C = 1$ and ridge $\alpha = 10$ were fixed. No external score or follow-up label was used to select these parameters. Weak prediction of availability does not establish MCAR or MAR. Fivefold cross-fitting reduces use of each observation in its own fitted prediction; it does not repair missing confounders or selection models that omit important clinical variables.

#### S8. Source completion bounds and tipping points

Bounds condition on a defined hypothetical score for every baseline participant and the unrestricted $[0, 1]$ support. They are not confidence intervals. The missing-mean tipping point $u^*$ and shift $\delta^*/s_R$ quantify the unobserved-score assumption required for zero completed change. Positive shifts require unavailable scores to be higher than observed scores; negative shifts require lower scores.

| Visit / program | Full-support interval | $u^*$ | $\delta^*$ | $\delta^*/s_R$ |
|---|---|---|---|---|
| Discharge / Inflammatory | $[-0.0612, 0.0289]$ | 0.6794 | 0.0621 | 6.19 |
| Discharge / TNF/NF-kB | $[-0.0696, 0.0205]$ | 0.7723 | 0.1162 | 6.86 |
| Discharge / IFN-alpha | $[-0.0518, 0.0383]$ | 0.5747 | −0.1760 | −5.85 |
| Discharge / IFN-gamma | $[-0.0609, 0.0292]$ | 0.6762 | −0.0823 | −4.20 |
| Discharge / IL6/JAK/STAT3 | $[-0.0643, 0.0258]$ | 0.7140 | 0.0440 | 4.37 |
| Discharge / Complement | $[-0.0595, 0.0306]$ | 0.6609 | −0.0041 | −0.68 |
| Discharge / Coagulation | $[-0.0407, 0.0494]$ | 0.4514 | −0.0102 | −1.21 |
| M1 / Inflammatory | $[-0.0978, 0.0463]$ | 0.6788 | 0.0652 | 7.89 |
| M1 / TNF/NF-kB | $[-0.1084, 0.0357]$ | 0.7523 | 0.1002 | 7.16 |
| M1 / IFN-alpha | $[-0.0956, 0.0486]$ | 0.6630 | −0.0839 | −3.65 |
| M1 / IFN-gamma | $[-0.1051, 0.0391]$ | 0.7290 | −0.0257 | −1.60 |
| M1 / IL6/JAK/STAT3 | $[-0.1043, 0.0398]$ | 0.7239 | 0.0582 | 6.82 |
| M1 / Complement | $[-0.0985, 0.0457]$ | 0.6832 | 0.0217 | 3.79 |
| M1 / Coagulation | $[-0.0697, 0.0744]$ | 0.4837 | 0.0269 | 3.29 |
| M6 / Inflammatory | $[-0.1646, 0.0877]$ | 0.6525 | 0.0396 | 4.42 |
| M6 / TNF/NF-kB | $[-0.1789, 0.0734]$ | 0.7092 | 0.0570 | 3.98 |
| M6 / IFN-alpha | $[-0.1784, 0.0739]$ | 0.7072 | −0.0369 | −1.57 |
| M6 / IFN-gamma | $[-0.1881, 0.0641]$ | 0.7457 | −0.0071 | −0.47 |
| M6 / IL6/JAK/STAT3 | $[-0.1769, 0.0754]$ | 0.7013 | 0.0364 | 4.42 |
| M6 / Complement | $[-0.1703, 0.0820]$ | 0.6750 | 0.0139 | 2.19 |
| M6 / Coagulation | $[-0.1185, 0.1338]$ | 0.4697 | 0.0121 | 1.50 |

For the primary M6 inflammatory score, the unrestricted interval crosses zero but zero requires a missing mean 4.42 observed SD above the observed mean. The prespecified ±2-SD scenarios therefore retain a negative contrast. This coexistence of worst-case nonidentification and scale-specific robustness should be preserved in interpretation. All 81 scenario points per source visit/program are in `source_mnar_scenarios.csv`; their labels explicitly identify assumed, not observed, missing outcomes.

### S9. External marginal comparisons and conditional completion

These entries describe GSE123342 unlinked MI assays. D30 contains 64 assays and Y1 contains 37, compared with 65 admission assays. Pairwise change, patient bootstrap intervals, source-model validation, and authenticated retention fractions are not available. The bounds and tipping shifts in this table are conditional scenarios that additionally assume each later assay represents a distinct member of the baseline group. Public metadata do not verify that assumption.

| Visit / program | Marginal change | Conditional interval | $u^*$ | $\delta^*/s_R$ |
|---|---|---|---|---|
| D30 / Inflammatory | −0.0025 | $[-0.0117, 0.0036]$ | 0.7636 | 26.89 |
| D30 / TNF/NF-kB | −0.0055 | $[-0.0153, 0.0001]$ | 0.9944 | 54.37 |
| D30 / IFN-alpha | 0.0021 | $[-0.0097, 0.0057]$ | 0.6298 | −6.42 |
| D30 / IFN-gamma | 0.0030 | $[-0.0087, 0.0067]$ | 0.5674 | −12.12 |
| D30 / IL6/JAK/STAT3 | −0.0030 | $[-0.0136, 0.0018]$ | 0.8813 | 31.47 |
| D30 / Complement | −0.0010 | $[-0.0112, 0.0042]$ | 0.7256 | 11.86 |
| D30 / Coagulation | −0.0015 | $[-0.0084, 0.0070]$ | 0.5479 | 15.42 |
| Y1 / Inflammatory | −0.0046 | $[-0.2631, 0.1676]$ | 0.6109 | 1.70 |
| Y1 / TNF/NF-kB | −0.0082 | $[-0.2819, 0.1489]$ | 0.6544 | 2.27 |
| Y1 / IFN-alpha | 0.0039 | $[-0.3283, 0.1024]$ | 0.7622 | −0.46 |
| Y1 / IFN-gamma | 0.0044 | $[-0.3233, 0.1075]$ | 0.7504 | −0.74 |
| Y1 / IL6/JAK/STAT3 | −0.0065 | $[-0.3013, 0.1295]$ | 0.6994 | 2.05 |
| Y1 / Complement | −0.0032 | $[-0.2871, 0.1437]$ | 0.6665 | 1.55 |
| Y1 / Coagulation | −0.0049 | $[-0.1979, 0.2329]$ | 0.4594 | 1.94 |

PBMC and whole-blood measurements differ in cellular composition, and six months and one year are not matched horizons. A similar direction does not establish biological equivalence or individual prediction transfer. Under the stated subset scenario, the Y1 inflammatory contrast reaches zero with an unavailable-assay mean about 1.70 observed SD higher than the observed mean. This external scenario lies within the prespecified two-SD range, unlike the linked source primary endpoint. The comparison concerns assumption sensitivity across settings, not a formal between-study test.

### S10. Observable selection and estimator settings

The availability models predict whether a deposited follow-up assay exists. Their performance is not clinical risk prediction. Fold-specific intercept-only models use the training prevalence. Their pooled AUROC can deviate from 0.5 because fold prevalences differ; use 0.5 as the conceptual no-information discrimination reference and do not interpret this artifact as negative biological prediction.

| Visit / model | AUROC | Brier | Log loss | ESS | Max. weight |
|---|---|---|---|---|---|
| Discharge / intercept | 0.496 | 0.082 | 0.303 | 101.0 | 1.10 |
| Discharge / logistic | 0.476 | 0.092 | 0.341 | 97.1 | 3.18 |
| M1 / intercept | 0.475 | 0.124 | 0.413 | 95.0 | 1.17 |
| M1 / logistic | 0.426 | 0.136 | 0.466 | 93.4 | 1.99 |
| M6 / intercept | 0.471 | 0.189 | 0.565 | 83.0 | 1.34 |
| M6 / logistic | 0.602 | 0.188 | 0.572 | 72.2 | 5.70 |

For the logistic model, ESS and maximum weight use the primary clipped probabilities. Intercept weights are constant. Complete predicted probabilities, quintile calibration bins, and baseline standardized differences are supplied as CSV files. The 99 M6 label permutations are a pipeline diagnostic and cannot test MAR.

#### Range across eight prespecified settings

Settings change one component at a time: logistic $C = 0.1$ or 10; ridge $\alpha = 1$ or 100; three or four folds; no propensity clipping; or clipping to $[0.1, 0.9]$. All unmodified components retain their primary setting. Ranges summarize these eight estimates, not uncertainty or a tuned best model. The full 168-row setting-by-score table is `model_sensitivity.csv`.

| Visit / program | IPW range | G-comp range | AIPW range |
|---|---|---|---|
| Discharge / Inflammatory | $[-0.0056, -0.0055]$ | $[-0.0060, -0.0056]$ | $[-0.0056, -0.0053]$ |
| Discharge / TNF/NF-kB | $[-0.0105, -0.0104]$ | $[-0.0112, -0.0106]$ | $[-0.0105, -0.0100]$ |
| Discharge / IFN-alpha | $[0.0165, 0.0202]$ | $[0.0159, 0.0162]$ | $[0.0170, 0.0188]$ |
| Discharge / IFN-gamma | $[0.0078, 0.0098]$ | $[0.0073, 0.0076]$ | $[0.0081, 0.0092]$ |
| Discharge / IL6/JAK/STAT3 | $[-0.0044, -0.0040]$ | $[-0.0044, -0.0040]$ | $[-0.0042, -0.0038]$ |
| Discharge / Complement | $[0.0004, 0.0006]$ | $[0.0002, 0.0003]$ | $[0.0004, 0.0005]$ |
| Discharge / Coagulation | $[0.0001, 0.0008]$ | $[0.0008, 0.0009]$ | $[-0.0000, 0.0006]$ |
| M1 / Inflammatory | $[-0.0093, -0.0087]$ | $[-0.0094, -0.0093]$ | $[-0.0092, -0.0086]$ |
| M1 / TNF/NF-kB | $[-0.0143, -0.0134]$ | $[-0.0148, -0.0144]$ | $[-0.0141, -0.0131]$ |
| M1 / IFN-alpha | $[0.0122, 0.0138]$ | $[0.0112, 0.0120]$ | $[0.0124, 0.0144]$ |
| M1 / IFN-gamma | $[0.0038, 0.0052]$ | $[0.0031, 0.0037]$ | $[0.0040, 0.0057]$ |
| M1 / IL6/JAK/STAT3 | $[-0.0083, -0.0078]$ | $[-0.0085, -0.0084]$ | $[-0.0082, -0.0076]$ |
| M1 / Complement | $[-0.0030, -0.0027]$ | $[-0.0031, -0.0030]$ | $[-0.0030, -0.0026]$ |
| M1 / Coagulation | $[-0.0037, -0.0033]$ | $[-0.0037, -0.0036]$ | $[-0.0037, -0.0032]$ |
| M6 / Inflammatory | $[-0.0098, -0.0095]$ | $[-0.0099, -0.0094]$ | $[-0.0105, -0.0097]$ |
| M6 / TNF/NF-kB | $[-0.0149, -0.0140]$ | $[-0.0140, -0.0136]$ | $[-0.0155, -0.0143]$ |
| M6 / IFN-alpha | $[0.0105, 0.0121]$ | $[0.0101, 0.0108]$ | $[0.0079, 0.0110]$ |
| M6 / IFN-gamma | $[0.0025, 0.0034]$ | $[0.0023, 0.0027]$ | $[0.0010, 0.0029]$ |
| M6 / IL6/JAK/STAT3 | $[-0.0089, -0.0088]$ | $[-0.0092, -0.0088]$ | $[-0.0094, -0.0090]$ |
| M6 / Complement | $[-0.0035, -0.0032]$ | $[-0.0036, -0.0033]$ | $[-0.0041, -0.0032]$ |
| M6 / Coagulation | $[-0.0038, -0.0029]$ | $[-0.0029, -0.0027]$ | $[-0.0049, -0.0030]$ |

#### S11.1. Artificial-deletion performance across programs

These experiments delete real assays from the finite parent of 83 M6 completers, not the 28 unavailable assays. Each mechanism/fraction uses 50 masks of the same parent. Entries are bias / RMSE in score units. Positive bias exceeds the known parent mean change. Nominal fractions 0.10, 0.25, and 0.40 delete 8/83, 21/83, and 33/83 assays.

| Program | Deletion | Fraction | Observed case | IPW | AIPW |
|---|---|---|---|---|---|
| Inflammatory | Uniform | 0.10 | 0.0000 / 0.0003 | −0.0001 / 0.0003 | −0.0002 / 0.0007 |
| Inflammatory | Uniform | 0.25 | 0.0000 / 0.0006 | −0.0001 / 0.0006 | −0.0003 / 0.0008 |
| Inflammatory | Uniform | 0.40 | −0.0001 / 0.0007 | 0.0000 / 0.0008 | −0.0002 / 0.0010 |
| Inflammatory | Baseline | 0.10 | −0.0003 / 0.0004 | 0.0000 / 0.0005 | −0.0006 / 0.0014 |
| Inflammatory | Baseline | 0.25 | −0.0010 / 0.0011 | 0.0005 / 0.0014 | 0.0005 / 0.0015 |
| Inflammatory | Baseline | 0.40 | −0.0016 / 0.0017 | −0.0009 / 0.0016 | −0.0001 / 0.0017 |
| Inflammatory | Follow-up | 0.10 | −0.0013 / 0.0013 | −0.0012 / 0.0013 | −0.0015 / 0.0016 |
| Inflammatory | Follow-up | 0.25 | −0.0032 / 0.0032 | −0.0029 / 0.0029 | −0.0032 / 0.0032 |
| Inflammatory | Follow-up | 0.40 | −0.0048 / 0.0048 | −0.0042 / 0.0043 | −0.0045 / 0.0046 |
| TNF/NF-kB | Uniform | 0.10 | −0.0000 / 0.0004 | −0.0003 / 0.0006 | −0.0007 / 0.0016 |
| TNF/NF-kB | Uniform | 0.25 | −0.0000 / 0.0010 | −0.0002 / 0.0010 | −0.0007 / 0.0018 |
| TNF/NF-kB | Uniform | 0.40 | −0.0002 / 0.0011 | −0.0000 / 0.0012 | −0.0005 / 0.0017 |
| TNF/NF-kB | Baseline | 0.10 | −0.0004 / 0.0006 | −0.0002 / 0.0010 | −0.0013 / 0.0031 |
| TNF/NF-kB | Baseline | 0.25 | −0.0014 / 0.0015 | 0.0008 / 0.0025 | 0.0009 / 0.0031 |
| TNF/NF-kB | Baseline | 0.40 | −0.0023 / 0.0024 | −0.0011 / 0.0022 | 0.0002 / 0.0021 |
| TNF/NF-kB | Follow-up | 0.10 | −0.0017 / 0.0017 | −0.0018 / 0.0020 | −0.0023 / 0.0028 |
| TNF/NF-kB | Follow-up | 0.25 | −0.0041 / 0.0041 | −0.0041 / 0.0041 | −0.0044 / 0.0045 |
| TNF/NF-kB | Follow-up | 0.40 | −0.0061 / 0.0061 | −0.0059 / 0.0060 | −0.0067 / 0.0069 |

Uniform deletion assigns equal sampling weights. Baseline- and follow-up-dependent deletion use the primary inflammatory score at the indicated visit: weights are proportional to $\exp\{2\,\mathrm{clip}(z, -2, 2)\}$, followed by fixed-size weighted sampling without replacement. Follow-up-dependent deletion explicitly uses information unavailable to the fitted baseline models. No mechanism is claimed to describe the actually unavailable participants. All masks, parent indices, scores, truth, estimates, and errors are saved; adjusted estimators need not dominate the observed-case contrast.

#### S11.2. Artificial-deletion performance across programs

These experiments delete real assays from the finite parent of 83 M6 completers, not the 28 unavailable assays. Each mechanism/fraction uses 50 masks of the same parent. Entries are bias / RMSE in score units. Positive bias exceeds the known parent mean change. Nominal fractions 0.10, 0.25, and 0.40 delete 8/83, 21/83, and 33/83 assays.

| Program | Deletion | Fraction | Observed case | IPW | AIPW |
|---|---|---|---|---|---|
| IFN-alpha | Uniform | 0.10 | 0.0002 / 0.0007 | 0.0005 / 0.0011 | −0.0001 / 0.0013 |
| IFN-alpha | Uniform | 0.25 | 0.0001 / 0.0014 | 0.0001 / 0.0013 | −0.0005 / 0.0019 |
| IFN-alpha | Uniform | 0.40 | −0.0002 / 0.0020 | 0.0002 / 0.0022 | −0.0004 / 0.0031 |
| IFN-alpha | Baseline | 0.10 | −0.0007 / 0.0011 | 0.0007 / 0.0030 | −0.0012 / 0.0041 |
| IFN-alpha | Baseline | 0.25 | −0.0020 / 0.0024 | 0.0002 / 0.0042 | 0.0004 / 0.0045 |
| IFN-alpha | Baseline | 0.40 | −0.0021 / 0.0025 | −0.0031 / 0.0041 | −0.0024 / 0.0047 |
| IFN-alpha | Follow-up | 0.10 | −0.0026 / 0.0028 | −0.0021 / 0.0024 | −0.0032 / 0.0038 |
| IFN-alpha | Follow-up | 0.25 | −0.0052 / 0.0053 | −0.0041 / 0.0044 | −0.0060 / 0.0062 |
| IFN-alpha | Follow-up | 0.40 | −0.0069 / 0.0070 | −0.0053 / 0.0058 | −0.0080 / 0.0085 |
| IFN-gamma | Uniform | 0.10 | 0.0001 / 0.0005 | 0.0003 / 0.0006 | −0.0001 / 0.0009 |
| IFN-gamma | Uniform | 0.25 | 0.0001 / 0.0009 | −0.0000 / 0.0009 | −0.0003 / 0.0013 |
| IFN-gamma | Uniform | 0.40 | −0.0002 / 0.0012 | 0.0001 / 0.0015 | −0.0003 / 0.0020 |
| IFN-gamma | Baseline | 0.10 | −0.0005 / 0.0007 | 0.0004 / 0.0019 | −0.0008 / 0.0027 |
| IFN-gamma | Baseline | 0.25 | −0.0014 / 0.0016 | 0.0003 / 0.0028 | 0.0004 / 0.0029 |
| IFN-gamma | Baseline | 0.40 | −0.0016 / 0.0019 | −0.0019 / 0.0027 | −0.0011 / 0.0031 |
| IFN-gamma | Follow-up | 0.10 | −0.0018 / 0.0019 | −0.0016 / 0.0017 | −0.0022 / 0.0025 |
| IFN-gamma | Follow-up | 0.25 | −0.0039 / 0.0040 | −0.0032 / 0.0034 | −0.0041 / 0.0042 |
| IFN-gamma | Follow-up | 0.40 | −0.0053 / 0.0054 | −0.0044 / 0.0046 | −0.0055 / 0.0059 |

Uniform deletion assigns equal sampling weights. Baseline- and follow-up-dependent deletion use the primary inflammatory score at the indicated visit: weights are proportional to $\exp\{2\,\mathrm{clip}(z,-2,2)\}$, followed by fixed-size weighted sampling without replacement. Follow-up-dependent deletion explicitly uses information unavailable to the fitted baseline models. No mechanism is claimed to describe the actually unavailable participants. All masks, parent indices, scores, truth, estimates, and errors are saved; adjusted estimators need not dominate the observed-case contrast.

#### S11.3. Artificial-deletion performance across programs

These experiments delete real assays from the finite parent of 83 M6 completers, not the 28 unavailable assays. Each mechanism/fraction uses 50 masks of the same parent. Entries are bias / RMSE in score units. Positive bias exceeds the known parent mean change. Nominal fractions 0.10, 0.25, and 0.40 delete 8/83, 21/83, and 33/83 assays.

| Program | Deletion | Fraction | Observed case | IPW | AIPW |
|---|---|---|---|---|---|
| IL6/JAK/STAT3 | Uniform | 0.10 | −0.0000 / 0.0002 | −0.0000 / 0.0003 | −0.0002 / 0.0006 |
| IL6/JAK/STAT3 | Uniform | 0.25 | −0.0001 / 0.0005 | −0.0001 / 0.0005 | −0.0003 / 0.0007 |
| IL6/JAK/STAT3 | Uniform | 0.40 | 0.0000 / 0.0007 | 0.0002 / 0.0008 | −0.0000 / 0.0008 |
| IL6/JAK/STAT3 | Baseline | 0.10 | −0.0003 / 0.0004 | 0.0002 / 0.0004 | −0.0004 / 0.0011 |
| IL6/JAK/STAT3 | Baseline | 0.25 | −0.0009 / 0.0010 | 0.0004 / 0.0011 | 0.0005 / 0.0012 |
| IL6/JAK/STAT3 | Baseline | 0.40 | −0.0016 / 0.0016 | −0.0008 / 0.0013 | 0.0001 / 0.0015 |
| IL6/JAK/STAT3 | Follow-up | 0.10 | −0.0010 / 0.0010 | −0.0008 / 0.0008 | −0.0010 / 0.0012 |
| IL6/JAK/STAT3 | Follow-up | 0.25 | −0.0025 / 0.0025 | −0.0020 / 0.0020 | −0.0023 / 0.0023 |
| IL6/JAK/STAT3 | Follow-up | 0.40 | −0.0039 / 0.0039 | −0.0029 / 0.0030 | −0.0034 / 0.0035 |
| Complement | Uniform | 0.10 | 0.0000 / 0.0002 | −0.0001 / 0.0003 | −0.0002 / 0.0005 |
| Complement | Uniform | 0.25 | 0.0000 / 0.0004 | −0.0000 / 0.0004 | −0.0001 / 0.0004 |
| Complement | Uniform | 0.40 | −0.0000 / 0.0005 | 0.0000 / 0.0006 | −0.0001 / 0.0007 |
| Complement | Baseline | 0.10 | −0.0002 / 0.0003 | 0.0001 / 0.0004 | −0.0002 / 0.0007 |
| Complement | Baseline | 0.25 | −0.0005 / 0.0006 | 0.0004 / 0.0011 | 0.0004 / 0.0010 |
| Complement | Baseline | 0.40 | −0.0008 / 0.0008 | −0.0004 / 0.0010 | 0.0000 / 0.0012 |
| Complement | Follow-up | 0.10 | −0.0007 / 0.0007 | −0.0006 / 0.0007 | −0.0008 / 0.0009 |
| Complement | Follow-up | 0.25 | −0.0017 / 0.0017 | −0.0015 / 0.0015 | −0.0016 / 0.0017 |
| Complement | Follow-up | 0.40 | −0.0027 / 0.0027 | −0.0024 / 0.0024 | −0.0026 / 0.0027 |

Uniform deletion assigns equal sampling weights. Baseline- and follow-up-dependent deletion use the primary inflammatory score at the indicated visit: weights are proportional to $\exp\{2\,\mathrm{clip}(z,-2,2)\}$, followed by fixed-size weighted sampling without replacement. Follow-up-dependent deletion explicitly uses information unavailable to the fitted baseline models. No mechanism is claimed to describe the actually unavailable participants. All masks, parent indices, scores, truth, estimates, and errors are saved; adjusted estimators need not dominate the observed-case contrast.

#### S11.4. Artificial-deletion performance across programs

These experiments delete real assays from the finite parent of 83 M6 completers, not the 28 unavailable assays. Each mechanism/fraction uses 50 masks of the same parent. Entries are bias / RMSE in score units. Positive bias exceeds the known parent mean change. Nominal fractions 0.10, 0.25, and 0.40 delete 8/83, 21/83, and 33/83 assays.

| Program | Deletion | Fraction | Observed case | IPW | AIPW |
|---|---|---|---|---|---|
| Coagulation | Uniform | 0.10 | 0.0000 / 0.0003 | −0.0001 / 0.0005 | −0.0003 / 0.0009 |
| Coagulation | Uniform | 0.25 | −0.0000 / 0.0005 | −0.0002 / 0.0005 | −0.0003 / 0.0008 |
| Coagulation | Uniform | 0.40 | −0.0001 / 0.0006 | −0.0001 / 0.0007 | −0.0002 / 0.0010 |
| Coagulation | Baseline | 0.10 | −0.0002 / 0.0004 | −0.0001 / 0.0006 | −0.0005 / 0.0013 |
| Coagulation | Baseline | 0.25 | −0.0006 / 0.0007 | 0.0006 / 0.0013 | 0.0006 / 0.0012 |
| Coagulation | Baseline | 0.40 | −0.0010 / 0.0011 | −0.0001 / 0.0012 | 0.0002 / 0.0012 |
| Coagulation | Follow-up | 0.10 | −0.0008 / 0.0009 | −0.0008 / 0.0010 | −0.0010 / 0.0014 |
| Coagulation | Follow-up | 0.25 | −0.0019 / 0.0020 | −0.0019 / 0.0020 | −0.0021 / 0.0022 |
| Coagulation | Follow-up | 0.40 | −0.0028 / 0.0028 | −0.0028 / 0.0029 | −0.0028 / 0.0029 |

Uniform deletion assigns equal sampling weights. Baseline- and follow-up-dependent deletion use the primary inflammatory score at the indicated visit: weights are proportional to $\exp\{2\,\mathrm{clip}(z,-2,2)\}$, followed by fixed-size weighted sampling without replacement. Follow-up-dependent deletion explicitly uses information unavailable to the fitted baseline models. No mechanism is claimed to describe the actually unavailable participants. All masks, parent indices, scores, truth, estimates, and errors are saved; adjusted estimators need not dominate the observed-case contrast.

#### S12. Analysis chronology, reproduction, and audit boundaries

The internal protocol was frozen on 27 October 2026 at 07:39:23 UTC, before score/effect/model inspection but after literature, metadata, availability-count, and prior-project review. It is not preregistration. A pre-score clarification restricted the external dataset to unlinked descriptive and conditional analysis. Fixed gene sets were acquired after the freeze; analysis used seed 20261027.

Post-analysis review identified a deviation: planned "rank-weighted" masking used $\exp\{2\,\mathrm{clip}(z,-2,2)\}$ of standardized inflammatory scores, without a participant-rank transform. Mechanisms, fractions, repetitions, and seed were retained; the exact formula was not fully prespecified. The frozen protocol is unchanged; see `protocol/implementation_deviation_01.txt`. The added baseline-correlation display is separately labeled exploratory.

##### Reproduction sequence

Use Python 3.11.9 and `requirements.txt` in a fresh extracted copy to preserve historical outputs. Set `OPENBLAS_NUM_THREADS=1` and `OMP_NUM_THREADS=1`; no GPU is needed. From the project root, retain the frozen acquisition manifests and run:

```
python code/acquire.py GSE59867 GSE123342
python code/download_annotations.py
python code/acquire_gene_sets.py
python code/prepare.py
python code/analyze.py --stage scores
python code/analyze.py --stage core
python code/analyze.py --stage controls
python code/summarize_results.py
python code/plot_figures.py
python code/validate_results.py
python code/build_supplement.py
python code/assemble.py --compile
```

Acquisition verifies and reuses matching inputs, rejecting changed bytes without replacing expected hashes or acquisition dates. Current remote files may differ from the snapshot. Preparation verifies matrix/annotation hashes; analysis checks protocol and score inputs. With annotations already present, `download_annotations.py` also accepts `-offline`. The compact release contains derived scores/results, not full matrices; direct plotting needs only derived files, whereas expression/rank validation requires preparation.

PDF reproduction additionally needs `pdflatex`, `bibtex`, and packages declared in `manuscript/main.tex`. Manifests record versions, hashes, time, seed, and repetitions.

##### Independent numerical audit

Without importing the production analysis module, the validator recomputes ranks, direct U comparisons, participant alignment, estimates, bounds, nuisance fits, all bootstraps, and every deletion mask. The initial 2,457 checks passed, including the logged discharge failure. This verifies computational consistency, not MAR, source consent, clinical utility, or biological interpretation.

##### Evidence boundaries

The study does not validate treatment effects, clinical prognosis, autonomous agents, or human–AI collaboration. Source measurements, hypothetical completion, and artificial deletion remain separate. No unavailable outcome is asserted as observed. Missingness reasons, survival, sampling decisions, cell counts, and linked clinical outcomes would be needed to narrow the identification limits. Review and revision records are stored separately from scientific results.
