## Supplementary material for "Follow-up availability and the robustness of immune transcriptomic recovery after myocardial infarction": Manuscript LaTeX source files: Figure_1_evidence_structure.pdf

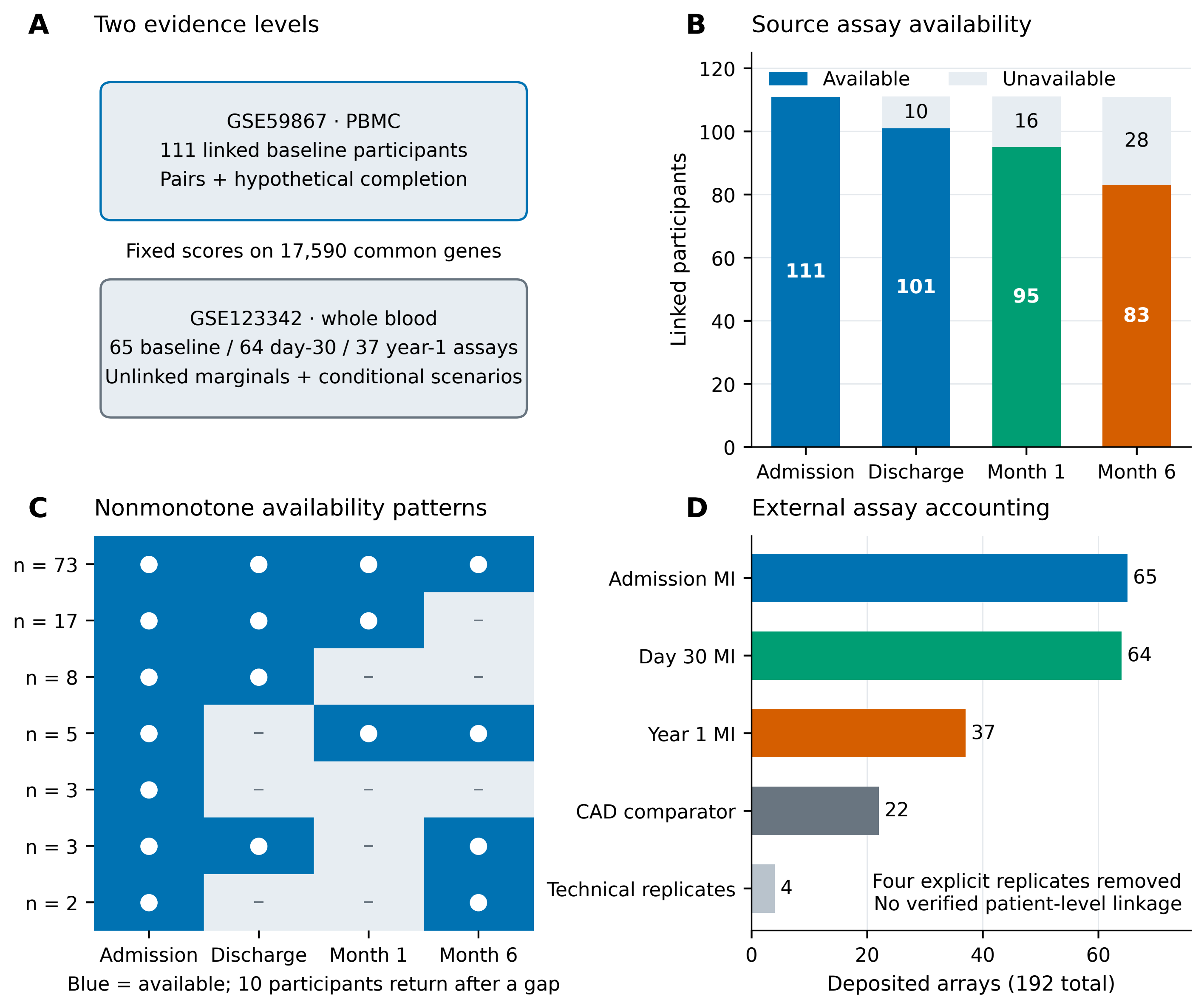
A
Two evidence levels
GSE59867 · PBMC
111 linked baseline participants
Pairs + hypothetical completion
Fixed scores on 17,590 common genes
GSE123342 · whole blood
65 baseline / 64 day-30 / 37 year-1 assays
Unlinked marginals + conditional scenarios
B
Source assay availability
Available
Unavailable
Linked participants
0
20
40
60
80
100
120
111
101
10
95
16
83
28
Admission
Discharge
Month 1
Month 6
C
Nonmonotone availability patterns
n = 73
n = 17
n = 8
n = 5
n = 3
n = 3
n = 2
Admission
Discharge
Month 1
Month 6
Blue = available; 10 participants return after a gap
D
External assay accounting
Admission MI
65
Day 30 MI
64
Year 1 MI
37
CAD comparator
22
Technical replicates
4
Four explicit replicates removed
No verified patient-level linkage
0
20
40
60
Deposited arrays (192 total)
