## Supplementary material for "Follow-up availability and the robustness of immune transcriptomic recovery after myocardial infarction": Manuscript LaTeX source files: Figure_2_observed_change.pdf

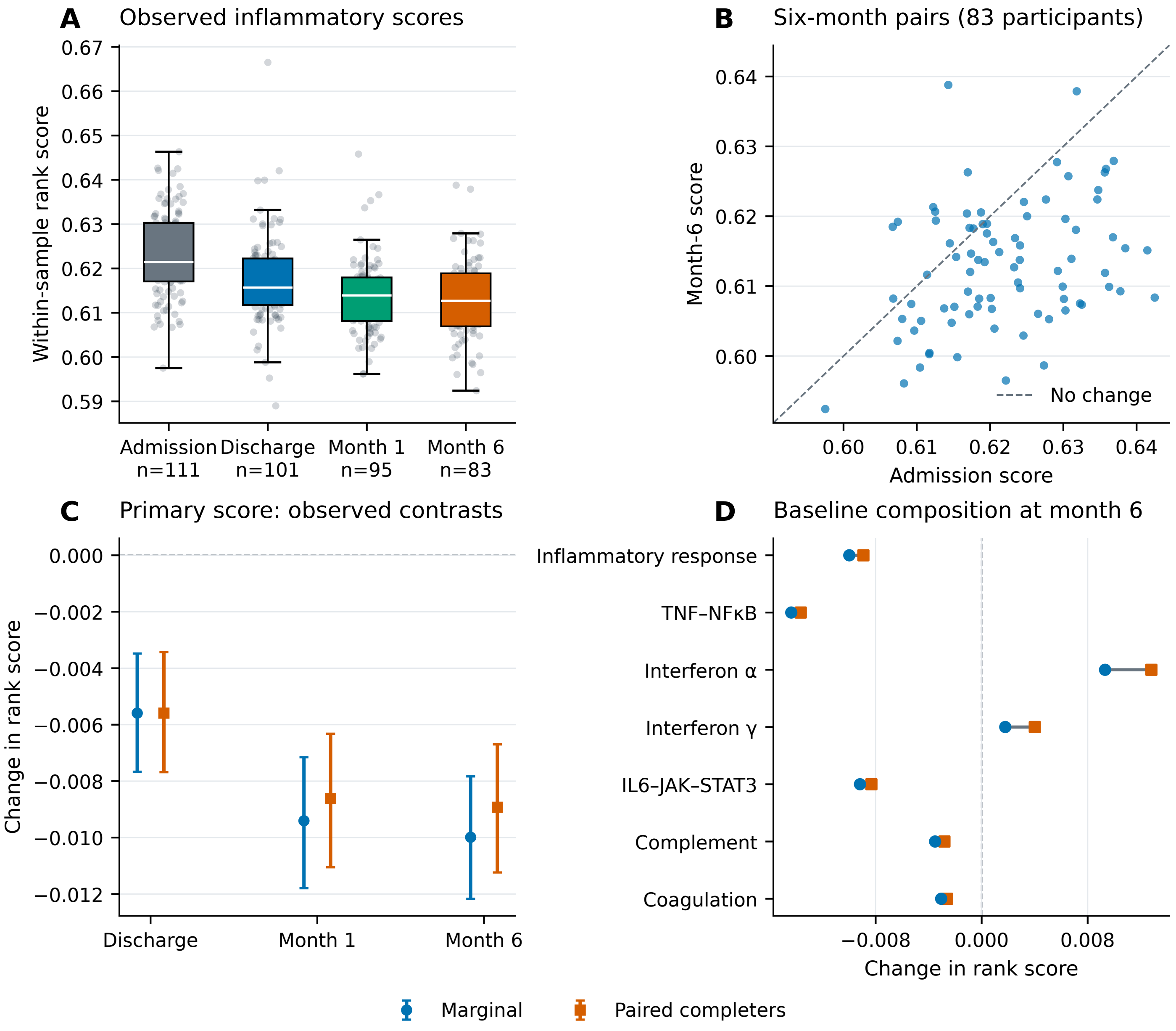
A
Observed inflammatory scores
Within-sample rank score
0.67
0.66
0.65
0.64
0.63
0.62
0.61
0.60
0.59
Admission n=111
Discharge n=101
Month 1 n=95
Month 6 n=83
B
Six-month pairs (83 participants)
Month-6 score
0.64
0.63
0.62
0.61
0.60
No change
0.60
0.61
0.62
0.63
0.64
Admission score
C
Primary score: observed contrasts
Change in rank score
0.000
−0.002
−0.004
−0.006
−0.008
−0.010
−0.012
Discharge
Month 1
Month 6
D
Baseline composition at month 6
Inflammatory response
TNF–NFκB
Interferon α
Interferon γ
IL6–JAK–STAT3
Complement
Coagulation
−0.008
0.000
0.008
Change in rank score
Marginal
Paired completers
