## Supplementary material for "Follow-up availability and the robustness of immune transcriptomic recovery after myocardial infarction": Manuscript LaTeX source files: Figure_3_identification_and_tipping.pdf

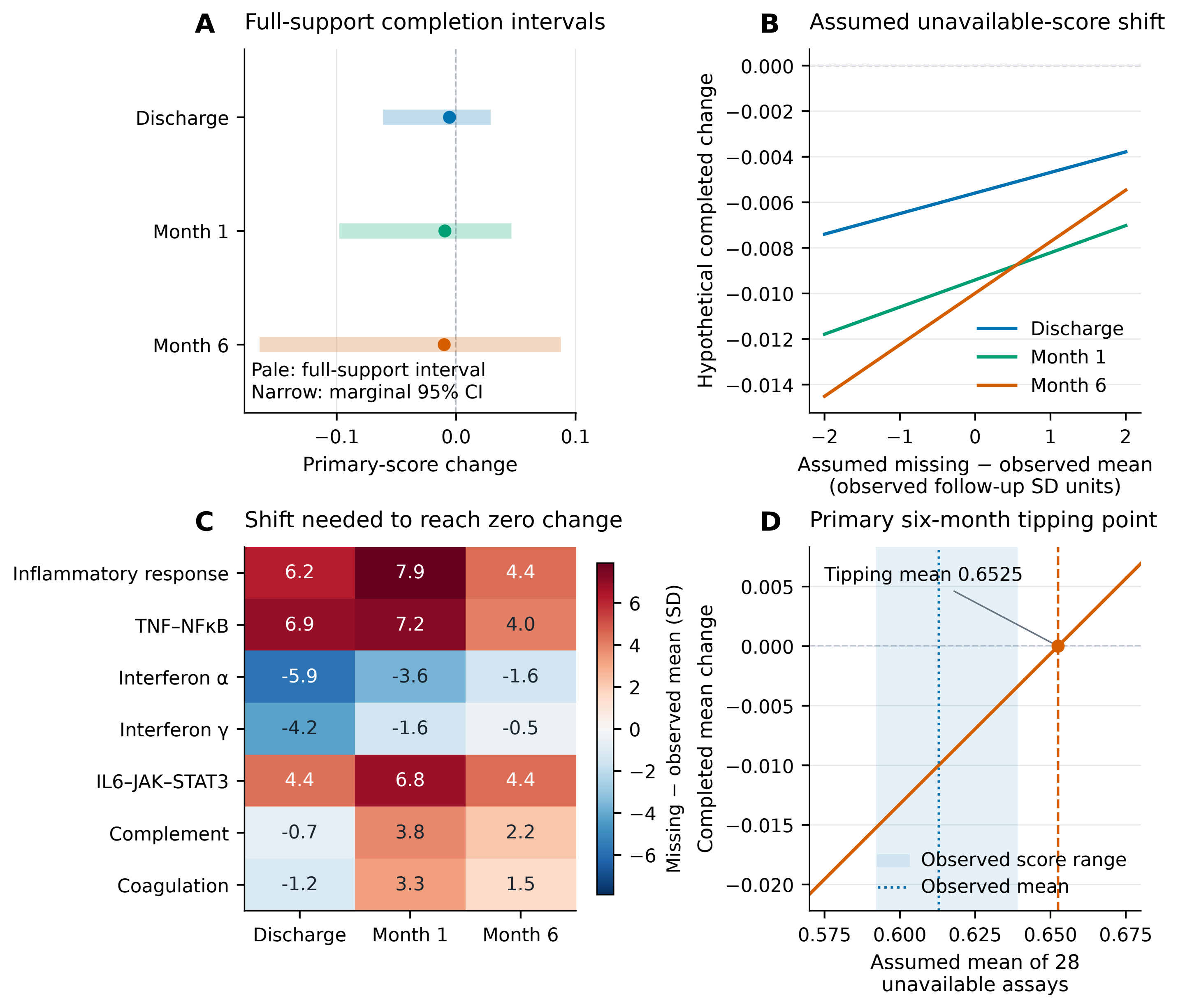
A
Full-support completion intervals
Discharge
Month 1
Month 6
Pale: full-support interval
Narrow: marginal 95% CI
−0.1
0.0
0.1
Primary-score change
B
Assumed unavailable-score shift
Hypothetical completed change
0.000
−0.002
−0.004
−0.006
−0.008
−0.010
−0.012
−0.014
Discharge
Month 1
Month 6
−2
−1
0
1
2
Assumed missing − observed mean
(observed follow-up SD units)
C
Shift needed to reach zero change
Inflammatory response
TNF–NFκB
Interferon α
Interferon γ
IL6–JAK–STAT3
Complement
Coagulation
6.2
7.9
4.4
6.9
7.2
4.0
-5.9
-3.6
-1.6
-4.2
-1.6
-0.5
4.4
6.8
4.4
-0.7
3.8
2.2
-1.2
3.3
1.5
Discharge
Month 1
Month 6
Missing − observed mean (SD)
6
4
2
0
−2
−4
−6
D
Primary six-month tipping point
Tipping mean 0.6525
Completed mean change
0.005
0.000
−0.005
−0.010
−0.015
−0.020
Observed score range
Observed mean
0.575
0.600
0.625
0.650
0.675
Assumed mean of 28
unavailable assays
