## Supplementary material for "Follow-up availability and the robustness of immune transcriptomic recovery after myocardial infarction": Manuscript LaTeX source files: Figure_4_availability_diagnostics.pdf

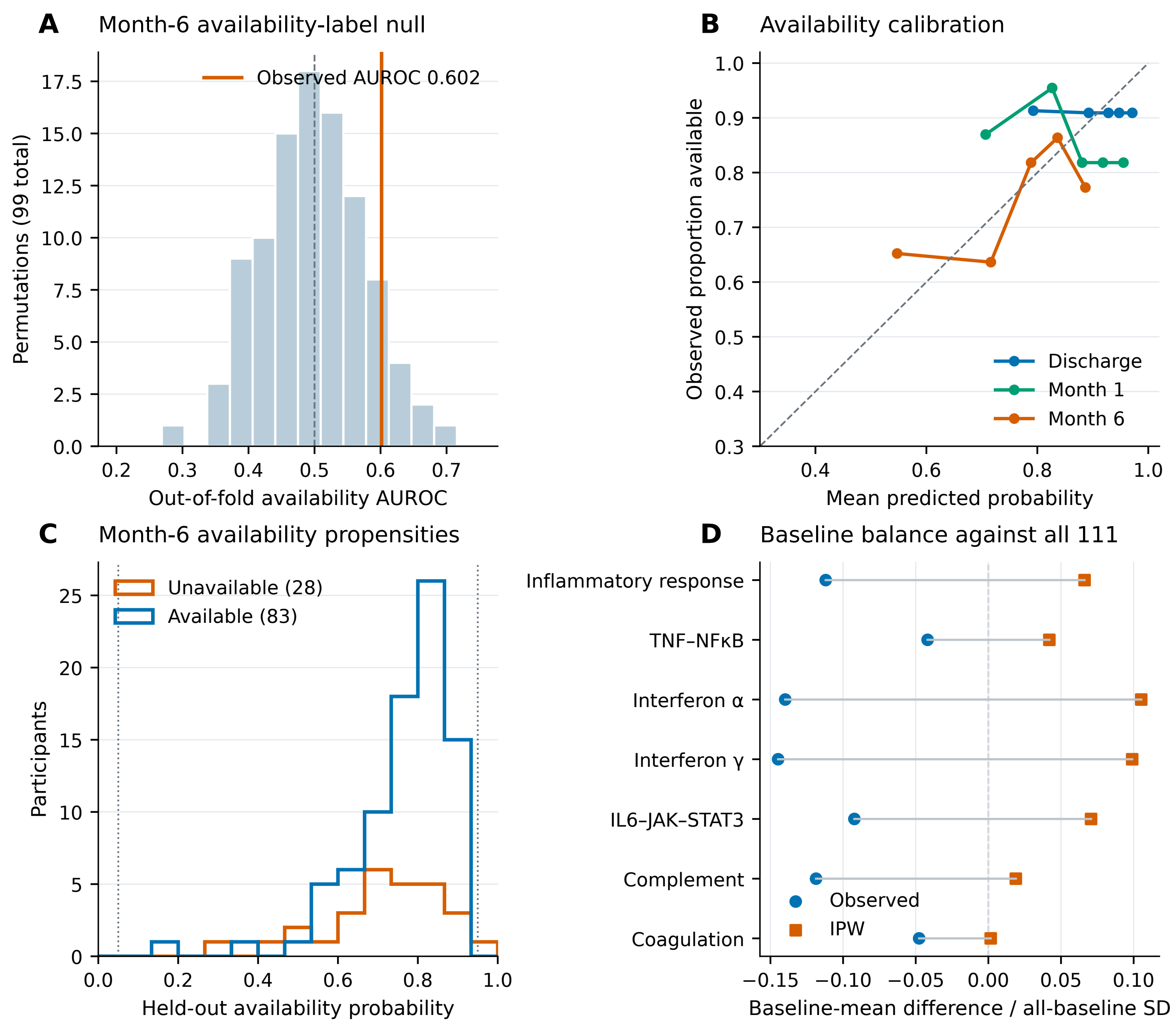
A
Month-6 availability-label null
Observed AUROC 0.602
Permutations (99 total)
Out-of-fold availability AUROC
B
Availability calibration
Observed proportion available
Mean predicted probability
Discharge
Month 1
Month 6
C
Month-6 availability propensities
Unavailable (28)
Available (83)
Participants
Held-out availability probability
D
Baseline balance against all 111
Inflammatory response
TNF–NFκB
Interferon α
Interferon γ
IL6–JAK–STAT3
Complement
Coagulation
Observed
IPW
Baseline-mean difference / all-baseline SD
