## Supplementary material for "Follow-up availability and the robustness of immune transcriptomic recovery after myocardial infarction": Manuscript LaTeX source files: Figure_5_mar_estimates.pdf

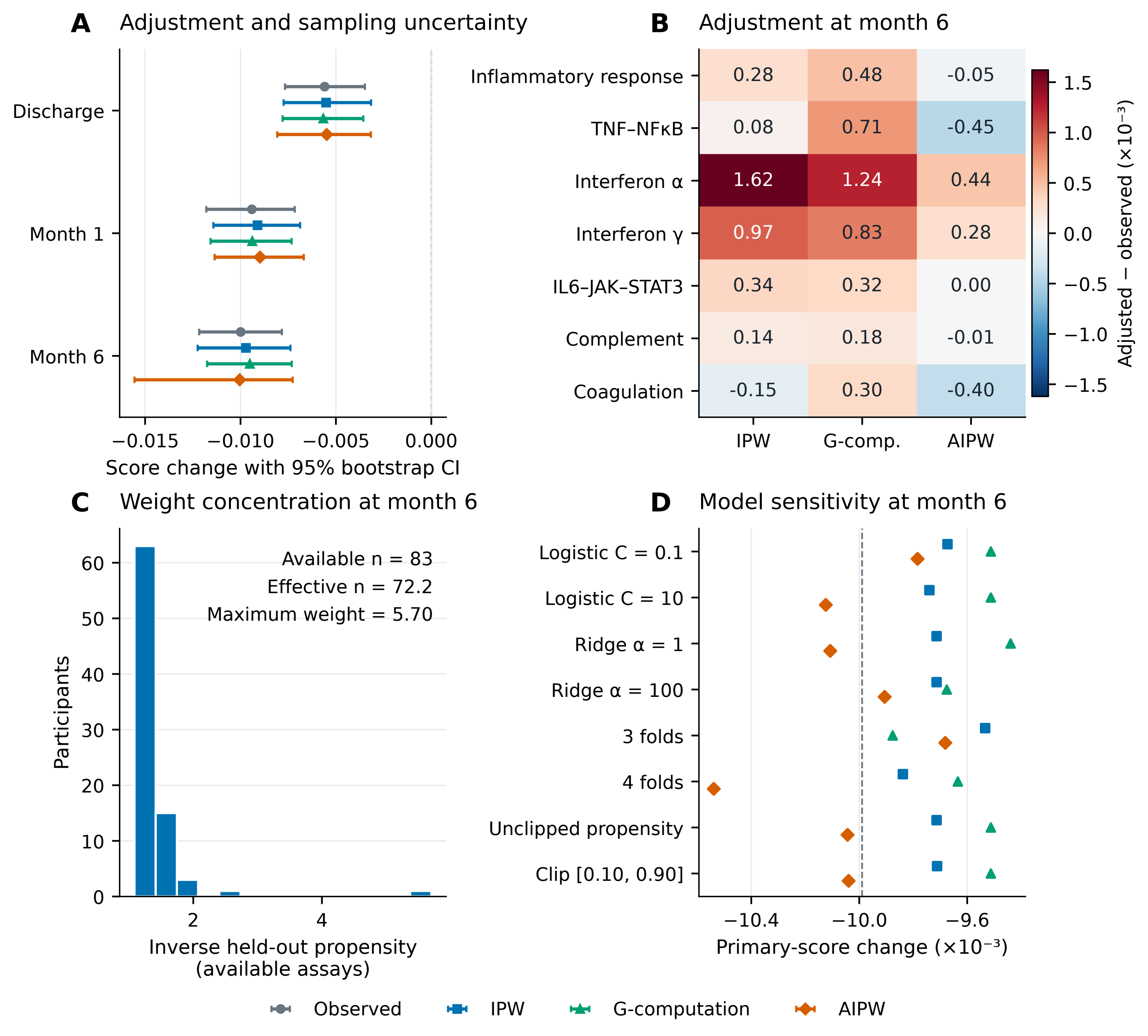

A
Adjustment and sampling uncertainty
Discharge
Month 1
Month 6
−0.015
−0.010
−0.005
0.000
Score change with 95% bootstrap CI
B
Adjustment at month 6
Inflammatory response
TNF–NFκB
Interferon α
Interferon γ
IL6–JAK–STAT3
Complement
Coagulation
0.28 0.48 -0.05
0.08 0.71 -0.45
1.62 1.24 0.44
0.97 0.83 0.28
0.34 0.32 0.00
0.14 0.18 -0.01
-0.15 0.30 -0.40
IPW
G-comp.
AIPW
Adjusted − observed (×10⁻³)
C
Weight concentration at month 6
Available n = 83
Effective n = 72.2
Maximum weight = 5.70
Participants
Inverse held-out propensity
(available assays)
D
Model sensitivity at month 6
Logistic C = 0.1
Logistic C = 10
Ridge α = 1
Ridge α = 100
3 folds
4 folds
Unclipped propensity
Clip [0.10, 0.90]
−10.4
−10.0
−9.6
Primary-score change (×10⁻³)
Observed
IPW
G-computation
AIPW
