## Supplementary material for "Follow-up availability and the robustness of immune transcriptomic recovery after myocardial infarction": Manuscript LaTeX source files: Figure_7_external_conditional_evidence.pdf

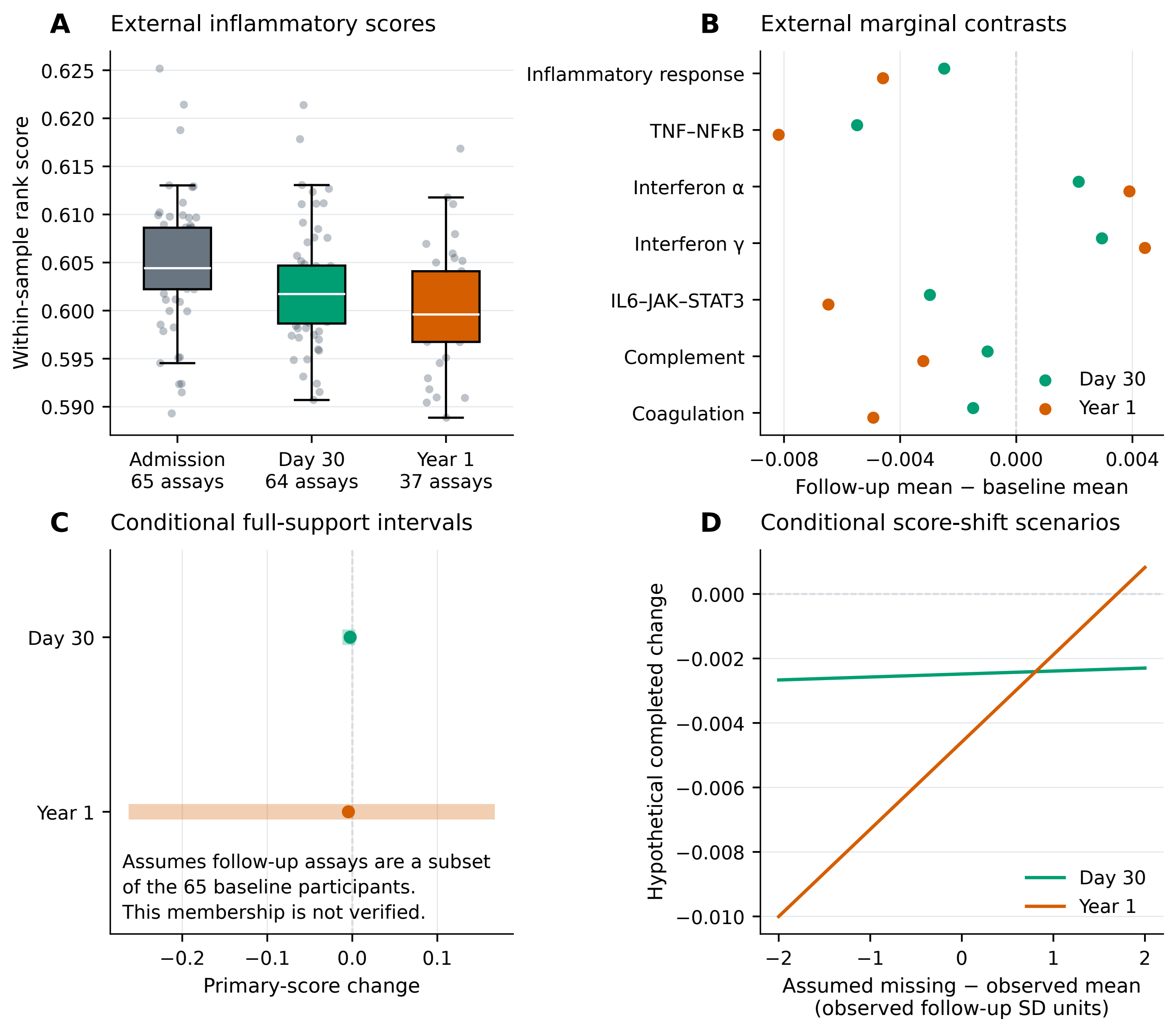
A
External inflammatory scores
Within-sample rank score
0.625
0.620
0.615
0.610
0.605
0.600
0.595
0.590
Admission
65 assays
Day 30
64 assays
Year 1
37 assays
B
External marginal contrasts
Inflammatory response
TNF–NFκB
Interferon α
Interferon γ
IL6–JAK–STAT3
Complement
Coagulation
Day 30
Year 1
−0.008
−0.004
0.000
0.004
Follow-up mean − baseline mean
C
Conditional full-support intervals
Day 30
Year 1
Assumes follow-up assays are a subset
of the 65 baseline participants.
This membership is not verified.
−0.2
−0.1
0.0
0.1
Primary-score change
D
Conditional score-shift scenarios
Hypothetical completed change
0.000
−0.002
−0.004
−0.006
−0.008
−0.010
Day 30
Year 1
−2
−1
0
1
2
Assumed missing − observed mean
(observed follow-up SD units)
