## Supplementary material for "Follow-up availability and the robustness of immune transcriptomic recovery after myocardial infarction": Manuscript LaTeX source files: Figure_8_score_definition_and_dependence.pdf

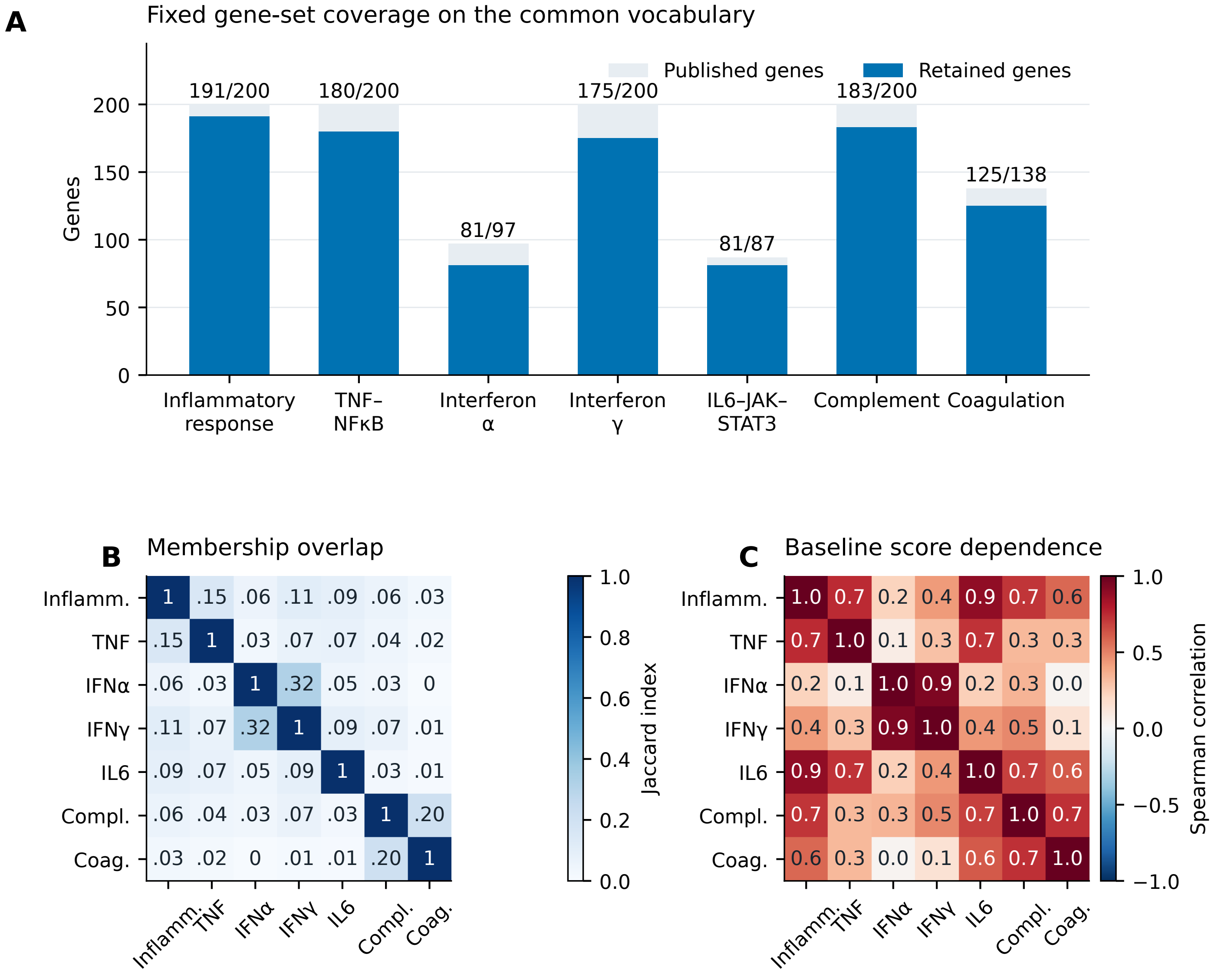
A
Fixed gene-set coverage on the common vocabulary
Published genes
Retained genes
Genes
191/200
180/200
81/97
175/200
81/87
183/200
125/138
Inflammatory response
TNF–NFκB
Interferon α
Interferon γ
IL6–JAK–STAT3
Complement
Coagulation
B
Membership overlap
Jaccard index
C
Baseline score dependence
Spearman correlation
Inflamm.
TNF
IFNα
IFNγ
IL6
Compl.
Coag.
