## Supplementary figures and images for "Follow-up availability and the robustness of immune transcriptomic recovery after myocardial infarction"

### Figure_1_evidence_structure.png

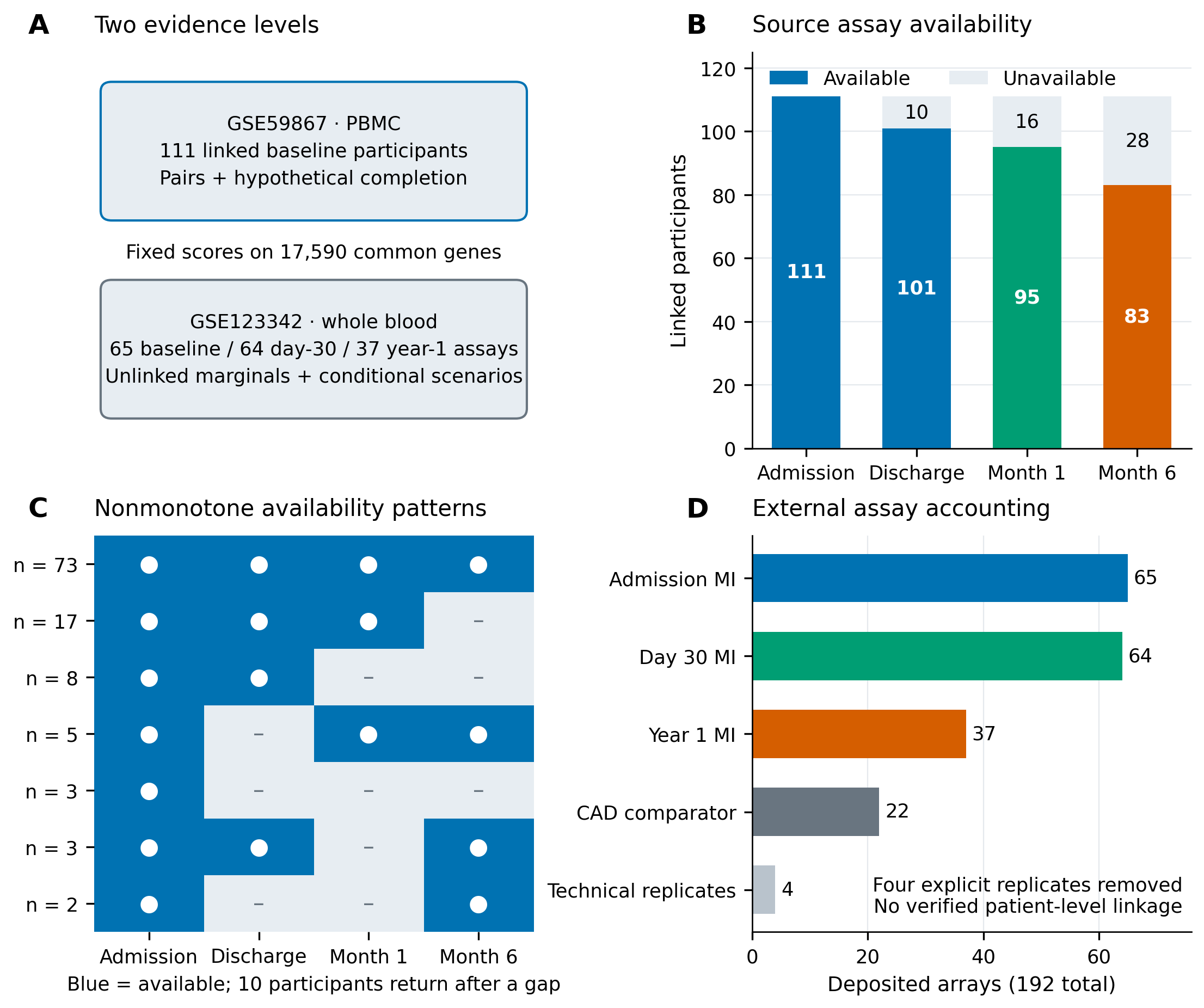

### Figure_2_observed_change.png

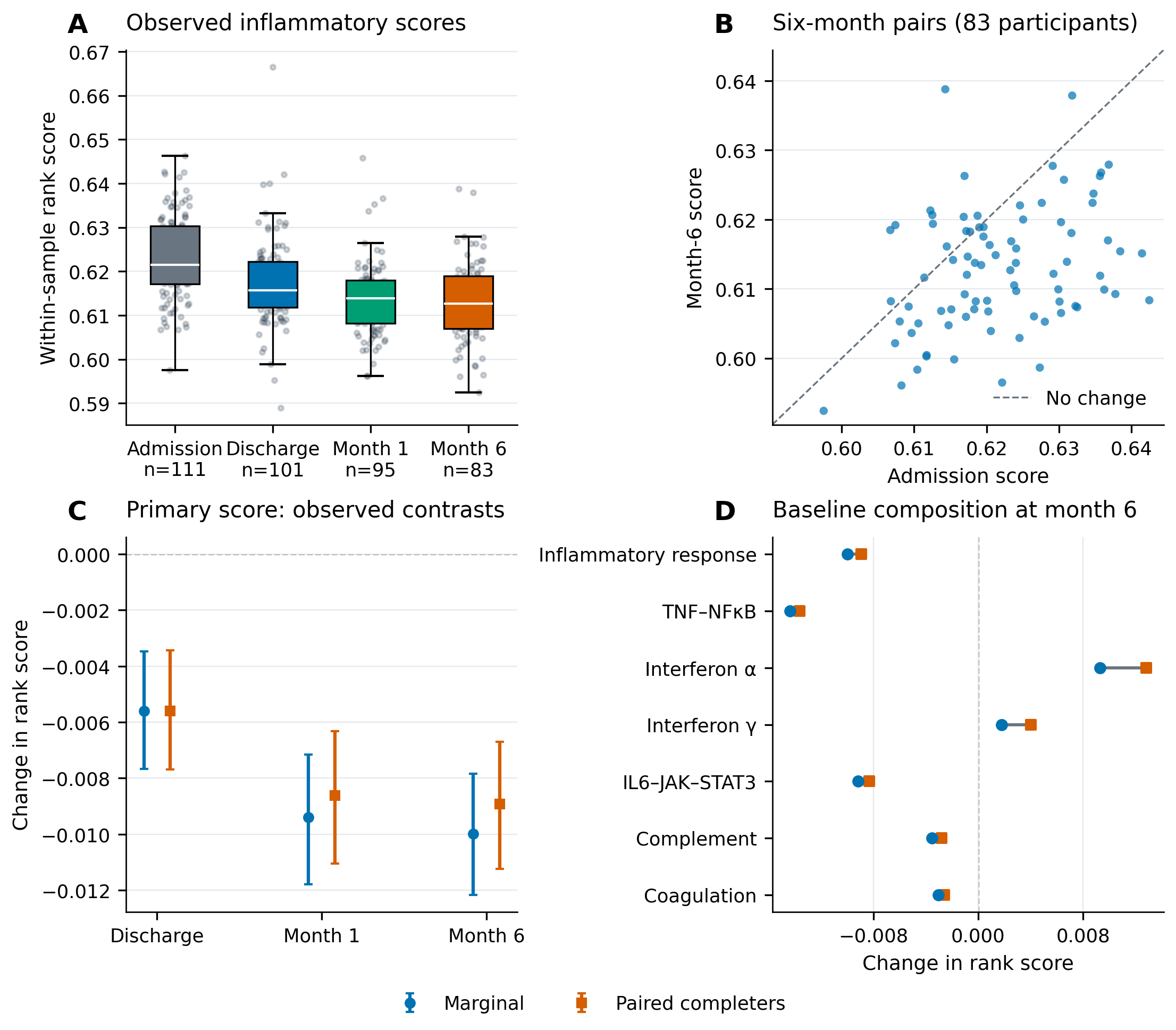

### Figure_3_identification_and_tipping.png

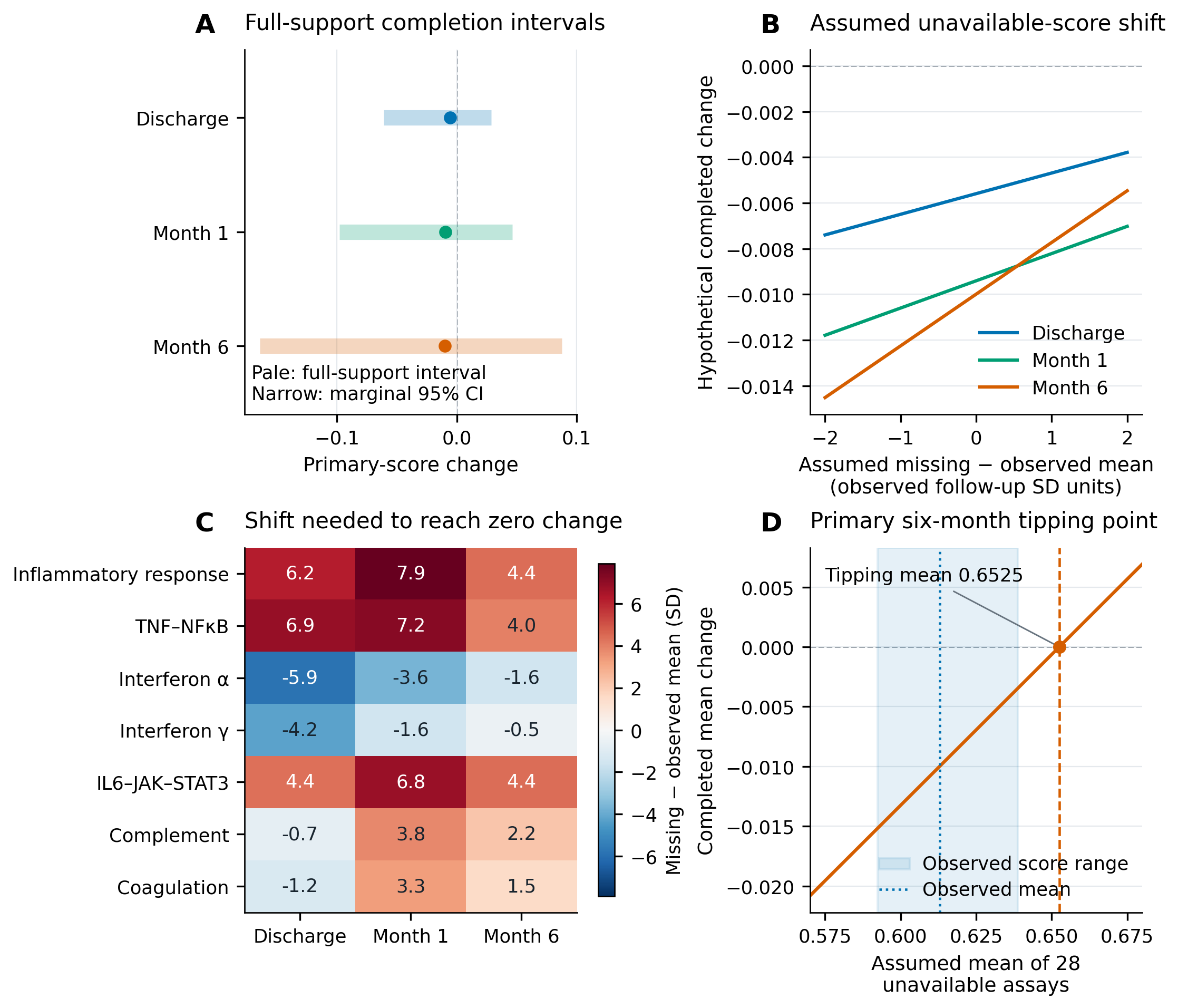

### Figure_4_availability_diagnostics.png

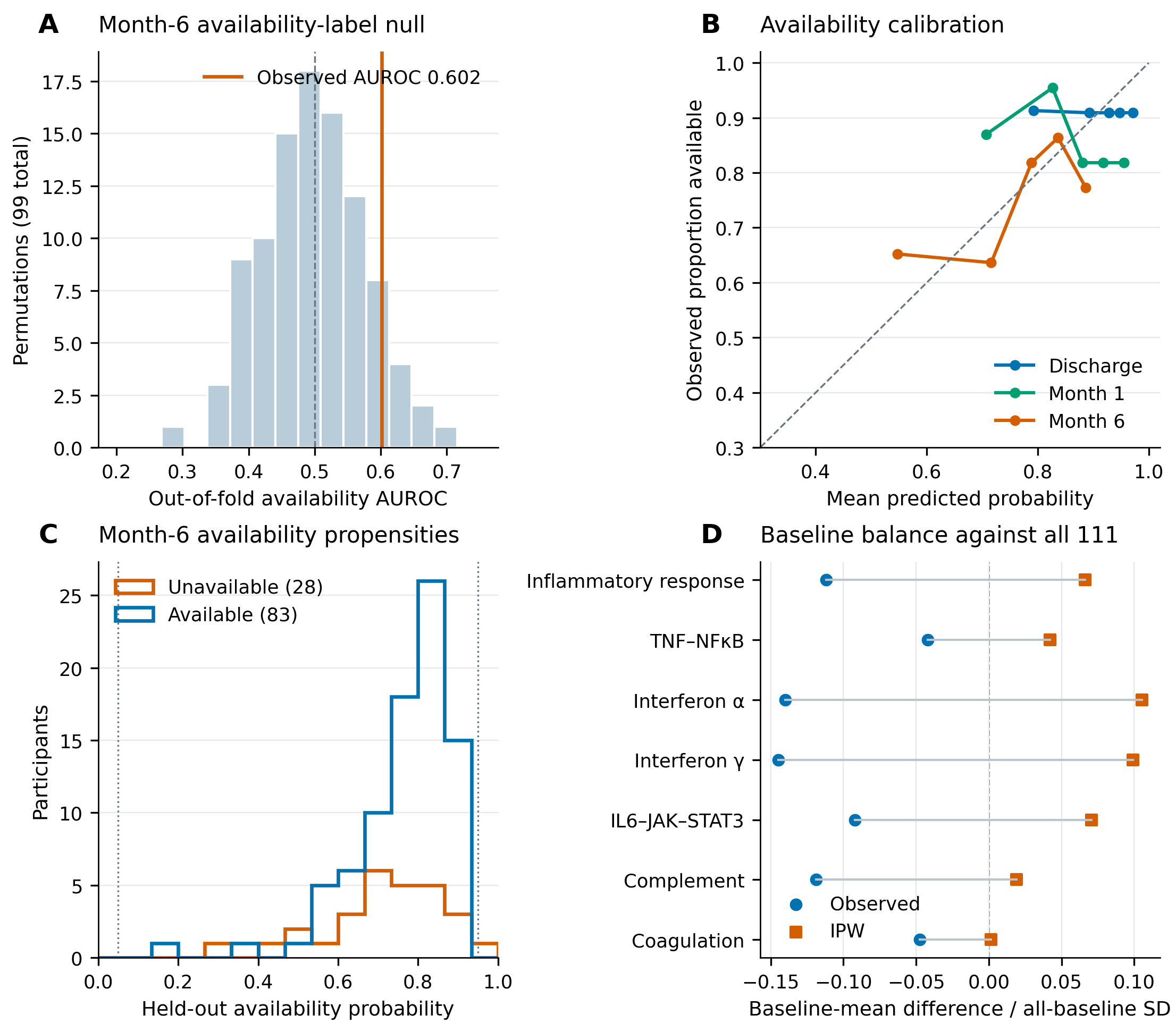

### Figure_5_mar_estimates.png

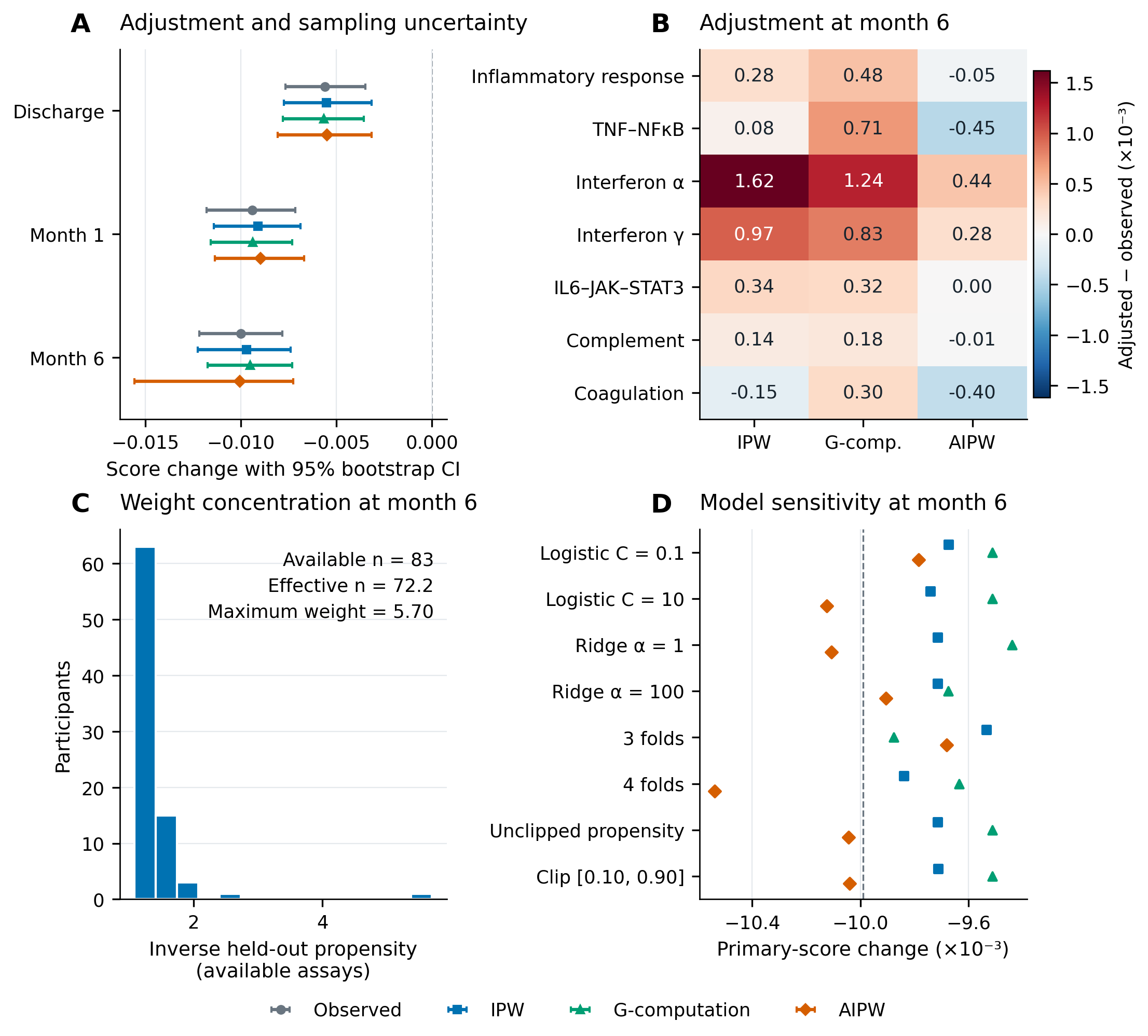

### Figure_6_artificial_deletion.pdf

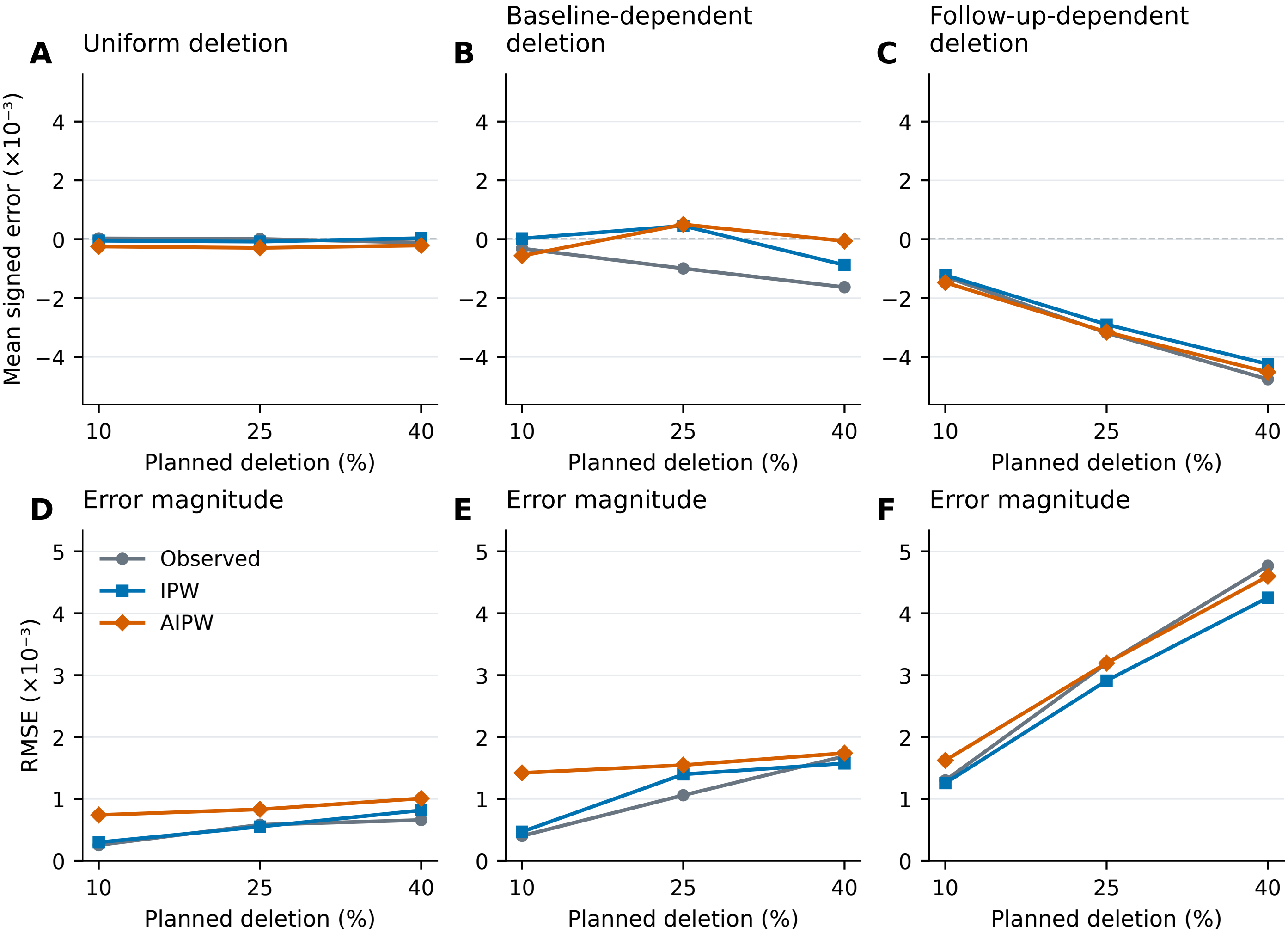

Artificial stress test: 83 observed completers; 50 masks per mechanism and fraction

### Figure_6_artificial_deletion.png

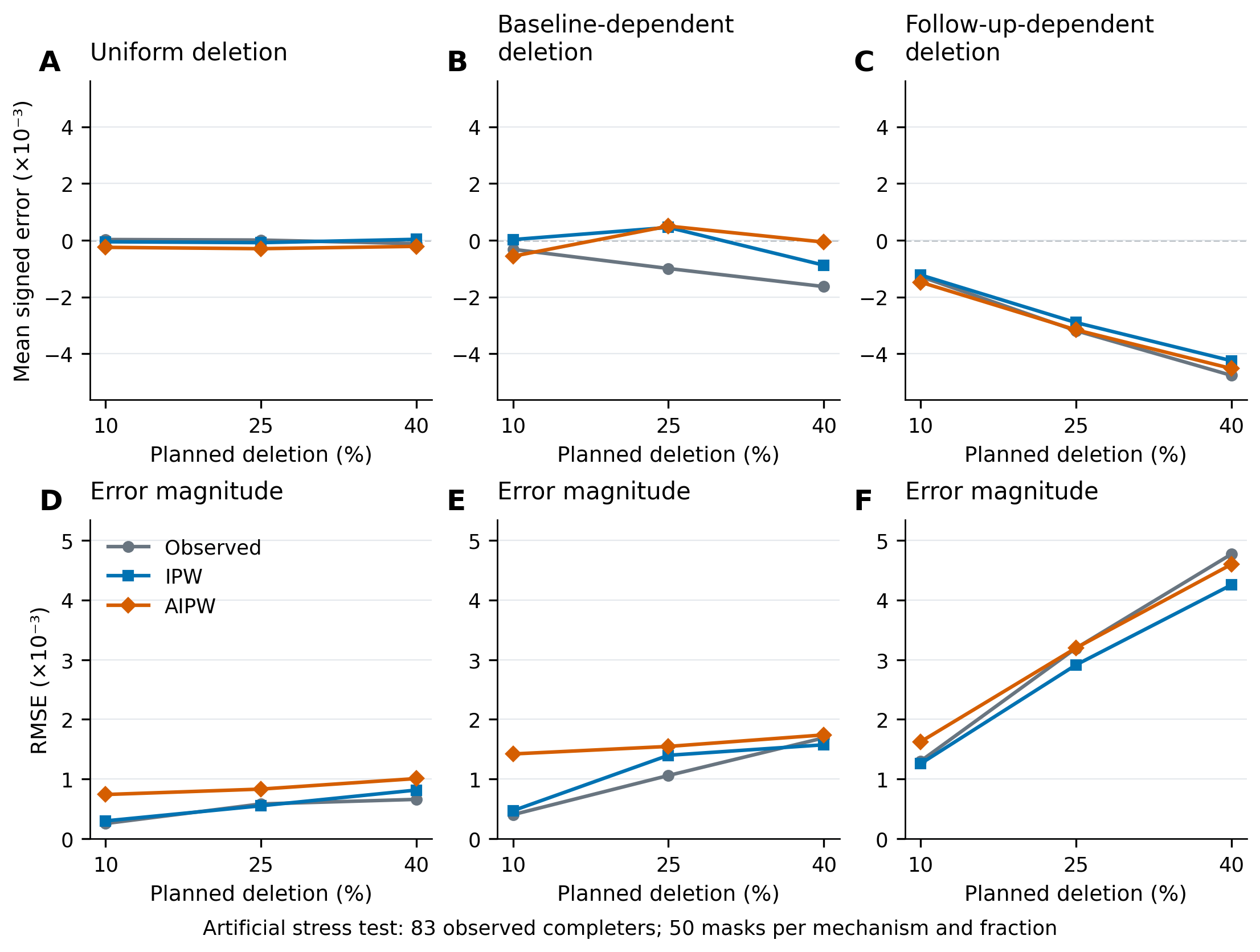

### Figure_7_external_conditional_evidence.png

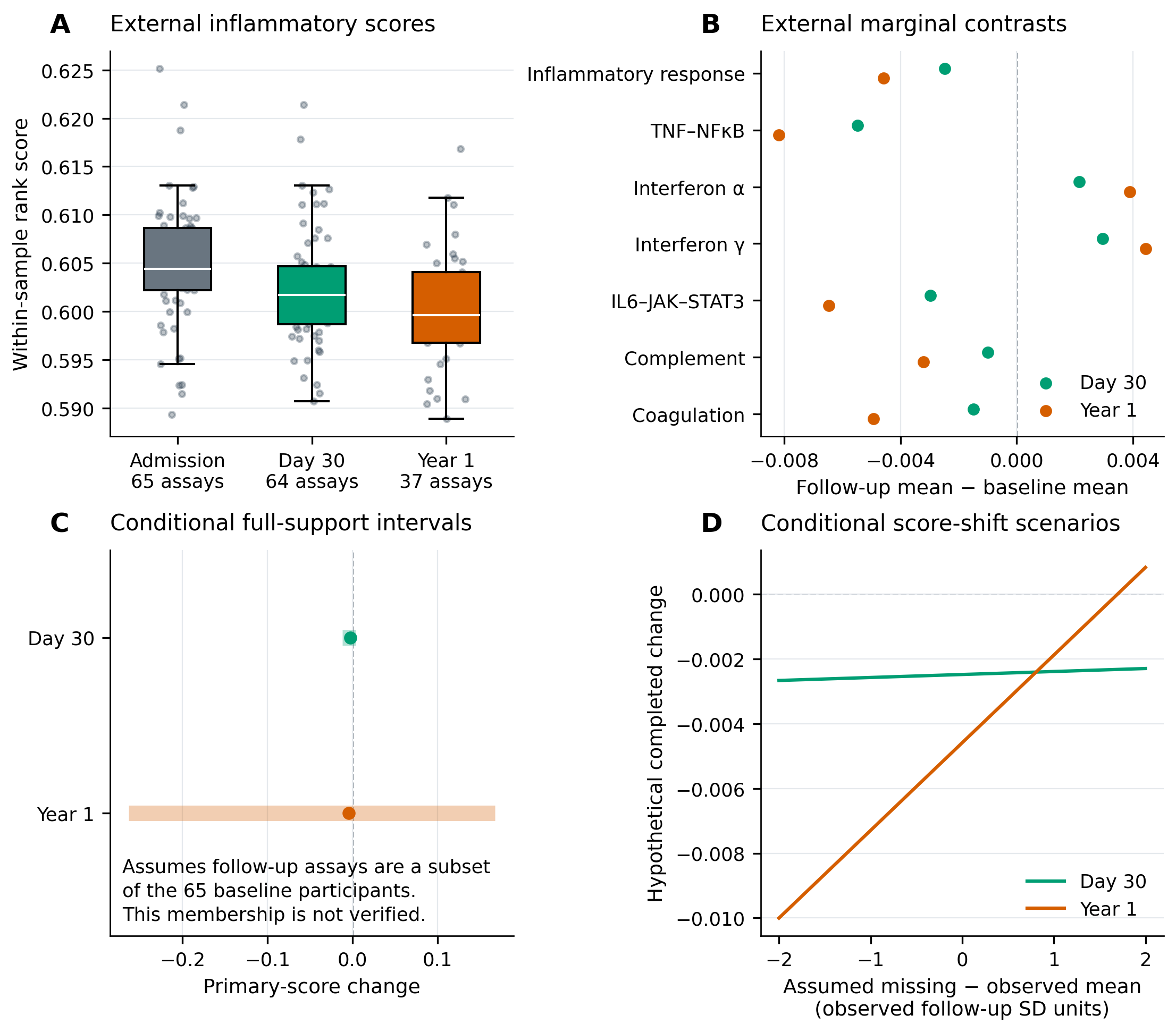

### Figure_8_score_definition_and_dependence.png

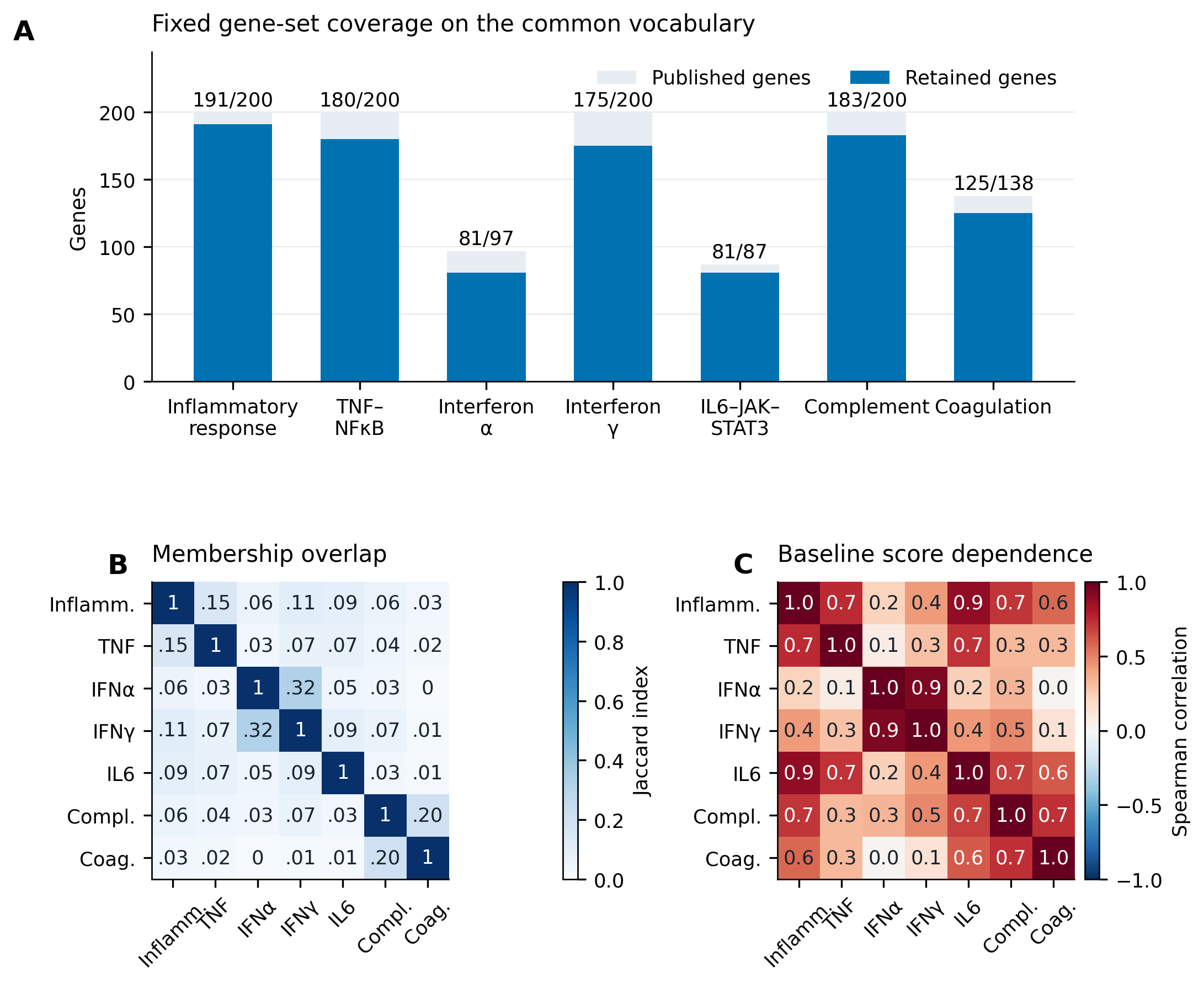
